# *Cis*-regulatory basis of sister cell type divergence in the vertebrate retina

**DOI:** 10.1101/648824

**Authors:** Daniel Murphy, Andrew. E.O. Hughes, Karen A. Lawrence, Connie A. Myers, Joseph C. Corbo

**Affiliations:** Deptartment of Pathology and Immunology, Washington University School of Medicine, St. Louis, MO; College of Social Work, University of Kentucky, Lexington, KY 40506

## Abstract

Multicellular organisms evolved via repeated functional divergence of transcriptionally related sister cell types, but the mechanisms underlying sister cell type divergence are not well understood. Here, we study a canonical pair of sister cell types, retinal photoreceptors and bipolar cells, to identify the key cis-regulatory features that distinguish them. By comparing open chromatin maps and transcriptomic profiles, we found that while photoreceptor and bipolar cells have divergent transcriptomes, they share remarkably similar cis-regulatory grammars, marked by enrichment of K50 homeodomain binding sites. However, cell class-specific enhancers are distinguished by enrichment of E-box motifs in bipolar cells, and Q50 homeodomain motifs in photoreceptors. We show that converting K50 motifs to Q50 motifs represses reporter expression in bipolar cells, while photoreceptor expression is maintained. These findings suggest that partitioning of Q50 motifs within cell type-specific cis-regulatory elements was a critical step in the divergence of the bipolar transcriptome from that of photoreceptors.

## Introduction

Complex tissues require the coordinated activity of a wide array of specialized cell types. It has been proposed that cell type diversity arises in the course of evolution through a ‘division of labor’ process, in which a multifunctional ancestral cell type gives rise to descendant cell types with divergent and novel functions^1,2^. Such descendants are often referred to as ‘sister’ cell types, and typically share a range of morphological, functional, and transcriptional features while at the same time displaying key differences^3,4^. A canonical example of sister cell types are mammalian retinal photoreceptors and bipolar cells^5,6^. In a typical vertebrate retina, photoreceptors synapse onto bipolar cells, which, in turn, synapse onto retinal ganglion cells that send their axons to the brain. Bipolar cells therefore constitute the central interneuronal cell class in the vertebrate retina. In mice, an array of 15 distinct bipolar cell types, broadly categorized as ON and OFF based on their response to light onset/offset, serve as a scaffold upon which the complex information-processing circuitry of the retina is built^7^. In this paper, we refer to photoreceptors and bipolar cells as cell ‘classes’, since they each comprise multiple distinct cell types.

During retinal development, photoreceptors and bipolar cells arise from the same population of OTX2-expressing progenitor cells^8–10^, share a similar elongate morphology^6^, and possess the molecular machinery required for ribbon synapse formation, a structure not found in any other retinal cell class^11^. In some vertebrate species, a subset of bipolar cells exhibit additional photoreceptor-like features, including localization of their cell bodies in the outer nuclear layer^12^ and the presence of an inner segment-like structure known as Landolt’s club, which extends from the dendrite to the outer limiting membrane and contains a 9+0 cilium^13–15^. These ‘transitional’ cell types point to the evolutionary origin of bipolar cells from photoreceptors^5,6^.

Both shared and divergent features of sister cell types are mediated by the transcriptional regulatory networks that govern gene expression in each cell type. In vertebrates, photoreceptors and bipolar cells express the paired-type homeodomain (HD) TFs CRX and OTX2, which are master regulators of gene expression in both cell classes^16–19^. A third paired-type HD TF, VSX2, is expressed specifically in bipolar cells and is required for bipolar fate^20,21^. Paired-type homeodomains recognize a core ‘TAAT’ motif, with additional specificity conferred by amino acids in positions 47, 50, and 54 of the homeodomain^22–24^ In particular, a lysine at position 50 (K50, as found in CRX and OTX2) favors recognition of TAAT<u>CC</u>, whereas a glutamine (Q50, as found in VSX2) favors recognition of TAATT^A^/_G_. Thus, substitution of a single amino acid in the HD toggles the TF’s binding preference for the nucleotides 3’ of the TAAT core^22^. Various bHLH TFs, which recognize E-box motifs (CANNTG), also play important roles in photoreceptor and bipolar cell gene expression programs. For instance, the bHLH TFs ASCL1 and NEUROD4 are required for the development of both photoreceptors and bipolar cells^25^. Similarly, NEUROD1 is required for photoreceptor survival, and BHLHE22 and BHLHE23 are required for development of OFF cone bipolar cells and rod bipolar cells, respectively^26–29^.

Our lab has previously shown that the *cis-*regulatory elements (CREs; i.e., enhancers and promoters) of mouse rods and cones are strongly enriched for K50 HD motifs as well as moderately enriched for Q50 HD and E-box motifs^30^. In addition, we recently used a massively parallel reporter assay to analyze the activity of thousands of photoreceptor enhancers identified by CRX ChIP-seq and found that both K50 HD and E-box motifs are positively correlated with enhancer activity in photoreceptors while Q50 HD motifs have a weakly negative correlation with enhancer activity^31^. In contrast, studies of individual reporters have shown that Q50 HD motifs mediate weak activation of expression via RAX, and that RAX can either enhance or suppress the transactivation activity of CRX, depending upon RAX expression levels^32,33^. Thus, Q50 HD motifs appear to have both positive and negative effects of photoreceptor enhancer activity, depending on context. In contrast, Q50 motifs in bipolar cells appear to be strongly repressive when bound by VSX2, which has been proposed to inhibit the expression of photoreceptor genes in bipolar cells^20,34,35^. The opposing effects on transcriptional activity mediated by K50 and Q50 motifs suggests that even subtle changes in HD binding sites may mediate major differences in gene expression. Indeed, a recent study in *Drosophila* showed that single base pair substitutions that interconvert Q50 and K50 half-sites within dimeric motifs are sufficient to switch the specificity of opsin expression within photoreceptor sub-types^36^.

Photoreceptors and bipolar cells offer an attractive model system in which to examine the mechanisms of *cis*-regulatory divergence in evolution and development, but the *cis*-regulatory landscape of bipolar cells is currently unknown. To elucidate the *cis*-regulatory grammar of bipolar cells we used FACS to isolate bipolar cell populations from mouse retina and obtain profiles of open chromatin and gene expression. By comparing these datasets to matching data from mouse rod and cone photoreceptors we found differential enrichment of Q50 motifs in photoreceptor-specific enhancers and a corresponding enrichment of E-boxes in bipolar-specific enhancers. We propose that the differential partitioning of Q50 motifs in photoreceptor and bipolar enhancers was a key evolutionary innovation contributing to transcriptomic divergence between the two cell classes.

## Results

### Photoreceptors and bipolar cells exhibit divergent transcriptional profiles

To obtain cell class-specific transcriptome profiles of mouse bipolar cells we used fluorescence-activated cell sorting (FACS) to purify bipolar cell populations from adult mice. We first isolated all bipolar cells using *Otx2*-GFP mice. This line harbors a GFP cassette knocked into the endogenous *Otx2* locus^37^. Adult *Otx2*-GFP mice display high-level GFP in bipolar cells and low-level expression in photoreceptors (Fig. 1B). To purify ON and OFF bipolar cells separately, we crossed *Otx2*-GFP mice with a *Grm6*-YFP line, in which YFP is driven by the *Grm6* promoter and expressed only in ON bipolar cells^38^. In the double transgenic mice (*Otx2*-GFP^+^; *Grm6*-YFP^+^) ON bipolar cells co-express GFP and YFP and can be separated from OFF bipolar cells based on fluorescence intensity (Fig. 1B). We subjected OFF bipolar cells to a second round of sorting to maximize purity from the adjacent weakly fluorescent photoreceptor population. Purity of bipolar cell populations was confirmed by qPCR which showed enrichment of ON- and OFF-specific genes in their respective populations and depletion of markers for other retinal cell classes as compared to whole retina (Fig. 1C). We then used RNA-seq to profile gene expression in purified populations of bipolar cells (Fig. S1).

**Figure 1.**
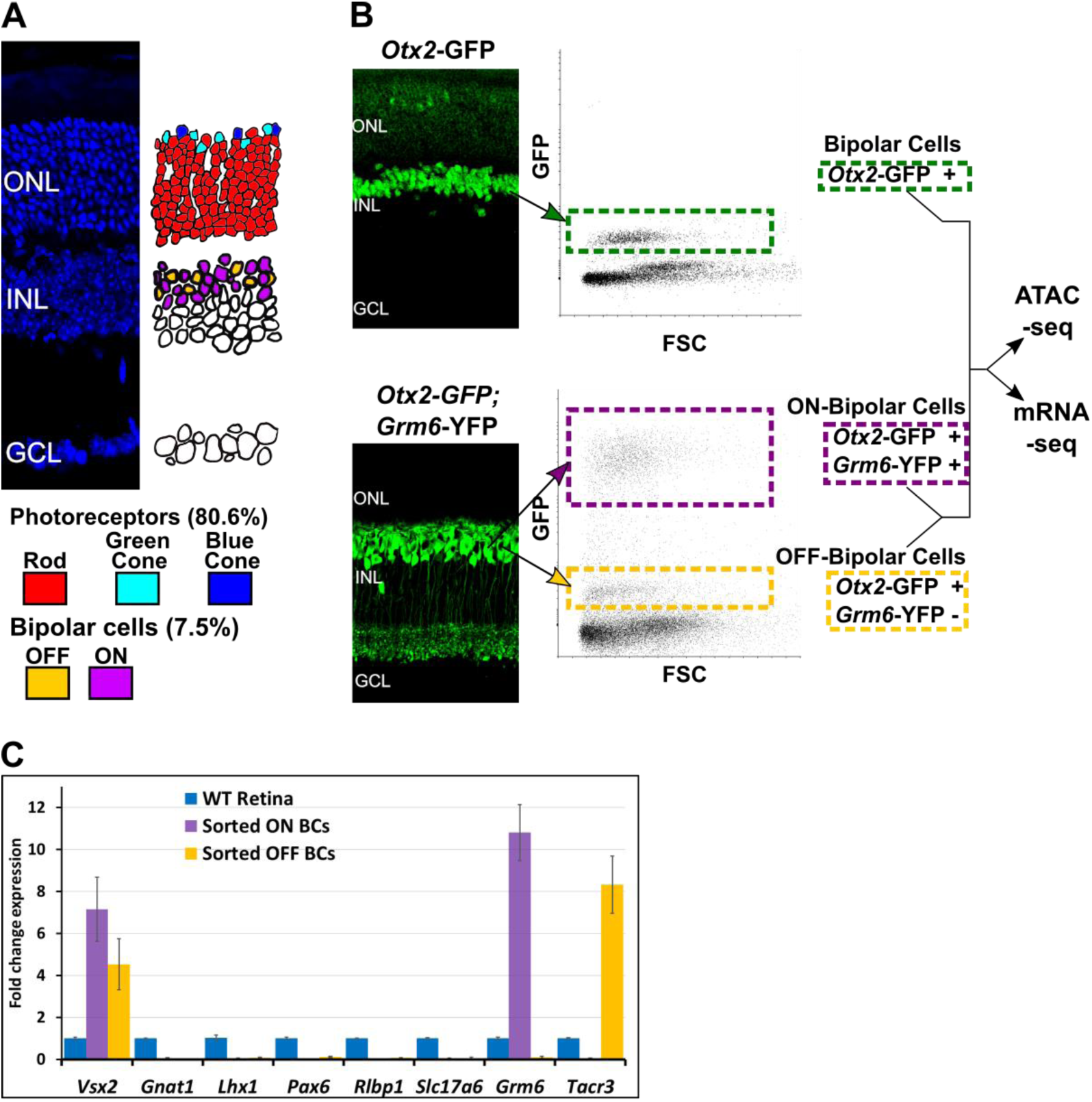
Isolation of bipolar cell populations from adult mouse retina. (**A**) **Left**, Histological section of adult mouse retina stained with 4′,6-diamidino-2-phenylindole (DAPI), to highlight nuclei. ONL= Outer Nuclear Layer, INL = Inner Nuclear Layer, GCL = Ganglion Cell Layer. **Right**, Schematic depiction of the location and relative abundance of photoreceptor and bipolar cell types. Percentage representation for each cell population in the mouse retina is shown, based on Jeon *et al.*^67^. (**B**) **Left**, Histologic sections of retina from transgenic mice expressing fluorescent marker proteins. In *Otx2*-GFP mice, GFP is strongly expressed in all bipolar cells, and weakly in photoreceptors. In *Grm6*-YFP mice, YFP is expressed exclusively in ON bipolar cells. Bipolar cell populations in the INL are linked to their position on FACS scatterplots with arrows (FSC= Forward Scatter). **Top**. All bipolar cells from *Otx2*-GFP^+^ mice are boxed in green. **Bottom**: in *Otx2*-GFP^+^;*Grm6*-YFP^+^ mice, ON bipolar cells (purple box) are separable from OFF bipolar cells (gold box) based on intensity of fluorescence. (**C**) RT-PCR analysis of retinal cell class markers^41^ in sorted ON (purple) and OFF (gold) bipolar cells normalized to expression in whole retina (blue). *Vsx2* = bipolar cells; *Gnat1* = rod photoreceptors; *Lhx1* = horizontal cells; *Pax6* = amacrine, ganglion, horizontal cells; *Rlbp1* = Müller glia; *Slc17a6* = ganglion cells; *Grm6* = ON bipolar cells; *Tacr3* = OFF bipolar cells (Types 1A, 1B, and 2).

To define the transcriptional differences between photoreceptor and bipolar cells we compared RNA-seq data from bipolar cells to similar data from wild-type rods and *Nrl*^−/−^ photoreceptors previously generated in our lab. We used *Nrl*^−/−^ photoreceptors as a surrogate for blue cones (i.e., *Opn1sw*-expressing cones), since mouse photoreceptors lacking *Nrl* transdifferentiate into blue cones during development^39,40^. We identified a total of 5,259 genes with at least a two-fold difference in expression between bipolar cells and either rods or blue cones (FDR < 0.05) (Table S5). Despite the large number of differentially expressed genes, published single-cell RNA-seq profiles indicate that the bipolar cell transcriptome is more similar to that of photoreceptors than to that of any other retinal cell class^41^. To evaluate the functional differences between photoreceptor and bipolar cell gene expression programs we compared the top ~30% most differentially expressed genes in each cell class (832 bipolar cell, 818 photoreceptor) using the gene ontology (GO) analysis tool, PANTHER^42^. Top bipolar-enriched GO terms were typical of many neuronal cell types and related to aspects of synaptic transmission, while photoreceptor-enriched GO terms mainly related to light-sensing (Table S6). To identify the transcriptional regulators, we compared the list of differentially expressed genes to a database of mouse TFs (AnimalTFDB3.0^43^), which revealed that 394 of the differentially expressed genes encode putative transcriptional regulators (Table S5). These include TFs known to be responsible for controlling gene expression in rods (*Nrl, Nr2e3,NeuroD1*), cones *(Thrb*), bipolar cells (*Vsx2*, *NeuroD4*), or both cell classes (*Crx*). Nearly one-third (176) of differentially expressed TFs are members of the zinc finger (ZF) family, many of which are more highly enriched in rods compared to bipolar cells but not in blue cone compared to bipolar cells. Conversely, of the top 10% differentially expressed TFs, the majority (25 of 35) are more highly expressed in bipolar cells compared to either rods or cones, and of these, most (16 of 25) are classified as HD, ZF or bHLH. Thus, the transcriptomes of bipolar cells and photoreceptors are surprisingly divergent, despite functional and morphological similarities between the two cell classes.

In contrast, comparison of the transcriptomes of ON and OFF bipolar cells identified only 680 genes that were differentially expressed by at least two-fold (317 ON- and 363 OFF-enriched at FDR < 0.05; Table S5). This figure is less than half of the number of differentially expressed genes identified between rods and blue cones (1,471), indicating that the transcriptomes of the two categories of bipolar cells are quite similar. A recent study by Shekhar *et al.*, described single-cell expression profiles for bipolar cell types using Drop-seq^44^. In order to verify our list of differentially expressed genes and gain insight into gene expression among individual bipolar cell types, we compared the list of genes differentially expressed between ON and OFF bipolar cells with the data of Shekhar *et al*. Overall, we found a strong correlation between the results of bulk ON and OFF bipolar cell expression profiling and single-cell transcriptome analysis (Fig. 2; Suppl. Fig. 2). However, we found that 50 of the 680 differentially expressed genes were not present in the Drop-seq data. These genes are generally expressed at low levels, even in the cell population in which they are enriched (Table S5). This result suggests that the greater sequence depth afforded by bulk RNA-seq allows for detection of subtle transcriptomic differences between cell populations that can be missed by single-cell profiling. Taken together, these data indicate that despite their sister cell type relationship, photoreceptor and bipolar cells have markedly distinct transcriptional profiles, while ON and OFF bipolar cells are more similar at the transcriptome level than rods and cones.

**Figure 2.**
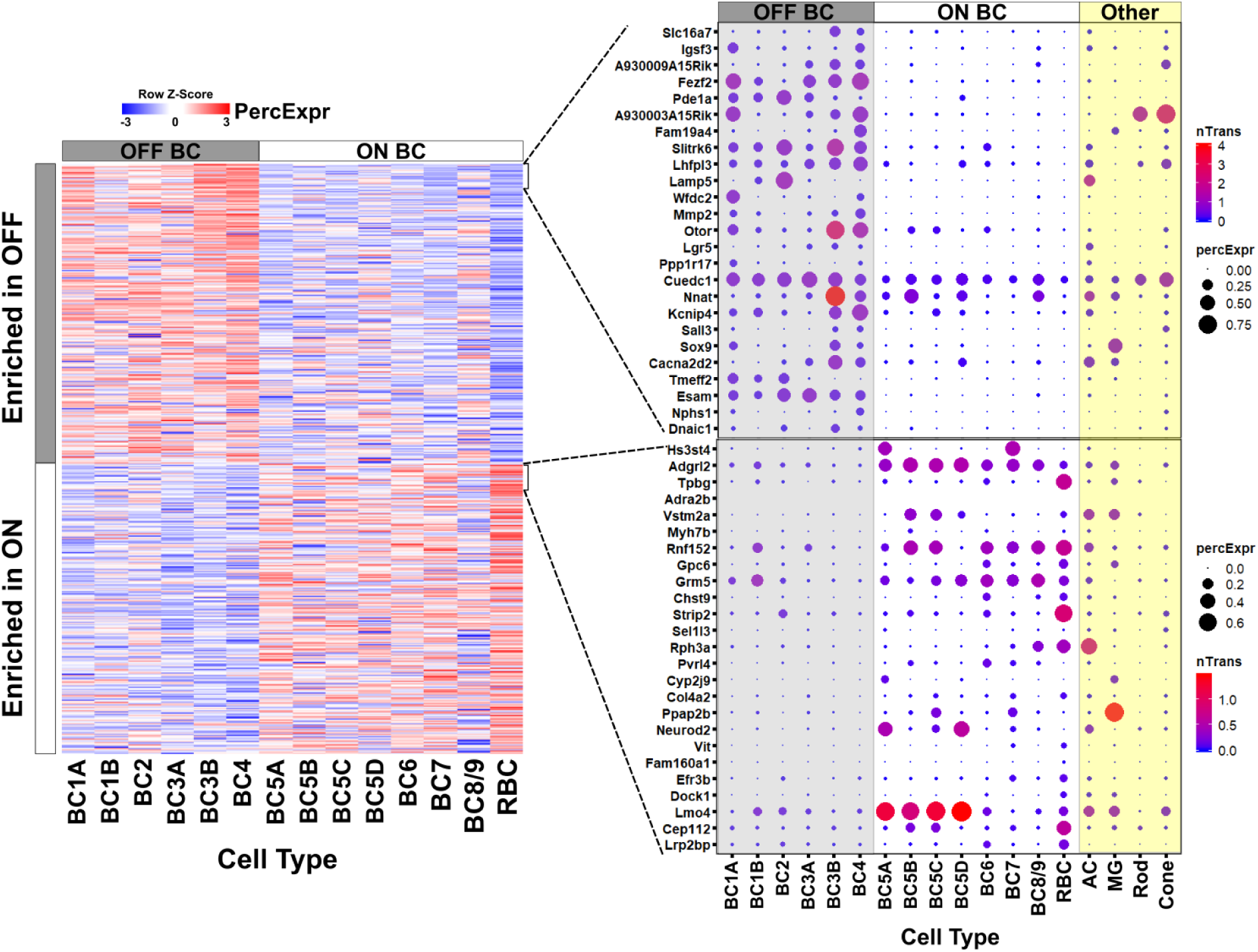
Gene expression in ON and OFF bipolar cells. **Left**, heatmap displaying 680 genes identified by bulk RNA-seq as differentially expressed between FACS-purified ON and OFF bipolar cell populations (current study) mapped onto single cell expression profiles for bipolar cell types identified by Drop-seq (Shekhar *et al.*)^44^. Overall, genes identified as ON- or OFF-specific by bulk RNA-seq showed corresponding differential expression between ON and OFF bipolar types identified by Dropseq. **Right**, Expression of the top 25 most differentially enriched genes (ranked by p-value) in OFF (top) and ON (bottom) bipolar populations presented as dot plots as in Shekhar *et al*. nTrans = mean number of transcripts expressed per cell in each cluster identified as a bipolar cell type; PercExpr = percentage of cells within each cluster found to express the indicated gene. While Drop-seq permits analysis of gene expression at single-cell resolution, bulk RNA-seq identified 50 differentially expressed genes not detected by Drop-seq (Table S5). Dot plots for all 680 differentially expressed genes are presented in Fig. S2.

### Bipolar cells have a more accessible chromatin landscape than either rods or cones

To compare chromatin accessibility between photoreceptor and bipolar cells, we used ATAC-seq (Assay for Transposase-Accessible Chromatin by sequencing) to generate open chromatin profiles from FACS-purified bipolar cells^45^. Similar to our RNA-seq data, ATAC-seq generated reproducible open chromatin profiles across biological replicates (Pearson correlation 0.95-1.00, Fig. S1). We combined ATAC-seq peaks from bipolar cells with previously generated ATAC-seq data from purified mouse rods, blue cones, and ‘green’ cones (i.e., *Opn1mw*-expressing cones) as well as DNAse-seq data^46^ from whole retina, brain, heart, and liver to obtain a list of >345,000 open chromatin regions. Hierarchical clustering of chromatin accessibility profiles at enhancers (i.e. regions >1000 bp upstream or >100 bp downstream of annotated transcription start sites) showed that photoreceptors, bipolar cells, and whole retina cluster separately from other tissues (Fig. 3A). Thus, the sister cell type relationship between photoreceptors and bipolar cells is reflected by the similarity of genome-wide patterns of enhancer chromatin accessibility. Interestingly, whole retina DNAse-seq clustered with bipolar cell samples, which is unexpected given that rod photoreceptors constitute ~80% of all cells in the mouse retina (Fig. 1A). This result may be a consequence of the distinctive pattern of global chromatin closure that we previously identified in rod photoreceptors^30^. Consistent with this, comparison of genome-wide patterns of chromatin accessibility in rods, cones and whole retina showed that more than half of the whole retina peaks did not overlap with photoreceptor peaks, suggesting that they likely derive from non-photoreceptor retinal cell types^30^.

**Figure 3.**
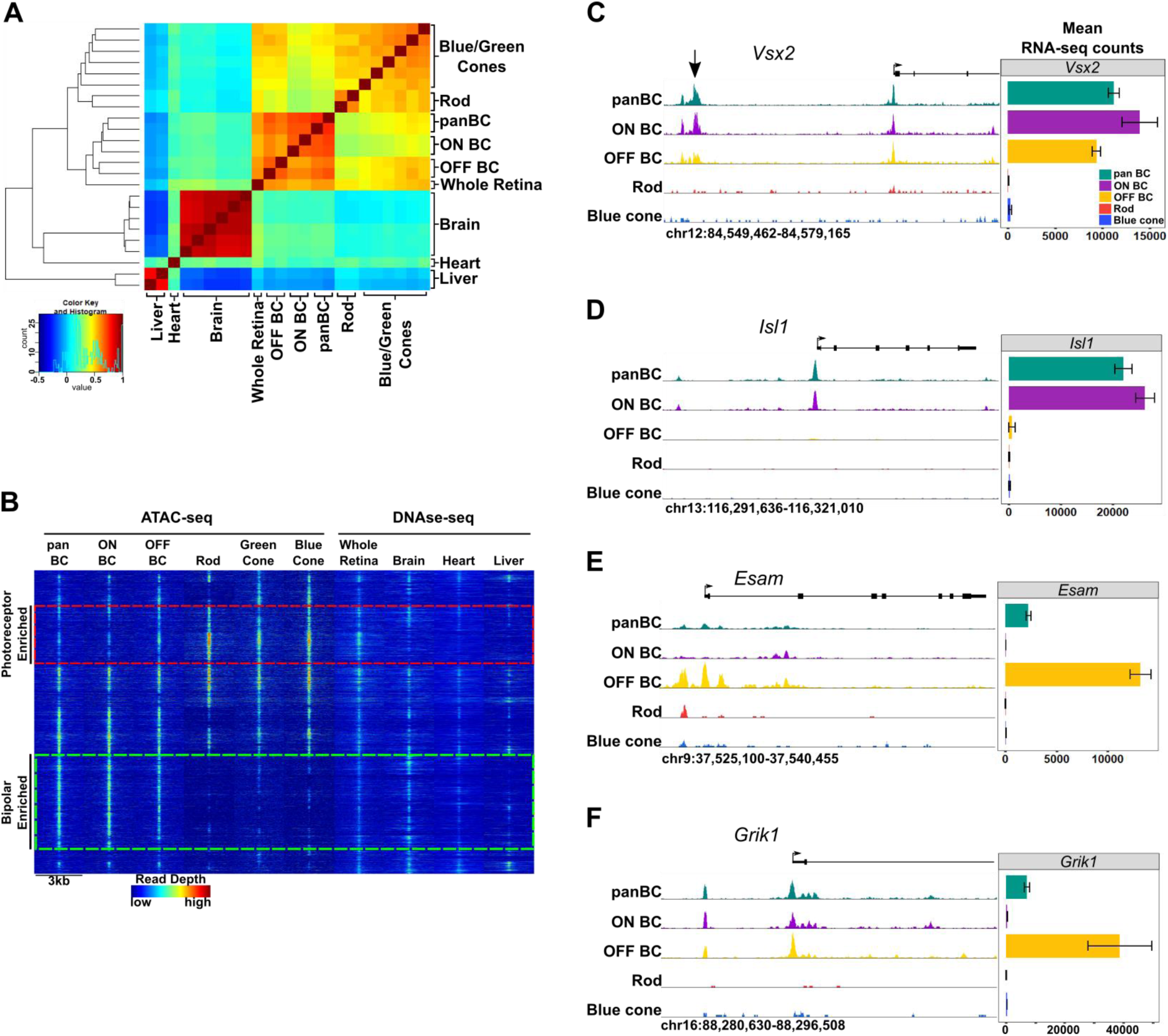
Genome-wide open chromatin profiles of ON and OFF bipolar cells. (**A**) Heatmap showing pairwise correlation between replicates of ATAC-seq data from photoreceptor and bipolar cell populations as well as DNAse-seq replicates from whole retina, brain, heart and liver. Peaks from each sample were combined to generate a set of 302,518 ‘enhancer’ (TSS-distal) peaks, and replicates were clustered based on read counts at each peak. Bipolar cells and photoreceptors form separate clusters. Whole retina DNAse-seq clusters with bipolar cells. Photoreceptors, bipolar cells and whole retina cluster separately from other tissues. (**B**) Genome-wide profiles of chromatin accessibility in isolated photoreceptor and bipolar cell ATAC-seq datasets as well as DNAse-seq datasets from additional control tissues. Rows show accessibility as indicated by read depth in 3 kb windows centered on peak summits sampled from photoreceptor, bipolar, and whole retina datasets (10,000 peaks randomly sampled from a total of 99,684 enhancer peaks are shown). Hierarchical clustering reveals peak sets enriched in photoreceptors (red box) or bipolar cells (green box). (**C-F**) Screenshots of UCSC genome browser tracks show regions of accessible chromatin in bipolar and photoreceptor populations at loci that exhibit shared or cell class-specific expression patterns. Black arrow in panel C indicates a known enhancer of *Vsx2*^54^. There is an imperfect correlation between chromatin accessibility and gene expression. Bar graphs aligned with browser tracks indicate mean RNA-seq counts of each gene for the indicated populations.

To investigate global accessibility within the retina we combined TSS-distal ATAC-seq peaks from bipolar cells and photoreceptors with DNAse-seq peaks from whole retina to create a list of 99,684 retinal open chromatin regions. Clustering these regions based on chromatin accessibility in retinal as well as non-retinal cell types offers a broad view of cell class- and cell type-specific regions of open chromatin (Fig. 3B). We found a large subset of bipolar-enriched peaks, many of which were also accessible in whole retina and brain (Fig. 3B, green box). Conversely, a smaller subset of peaks showed selective accessibility in photoreceptors, with lower levels of accessibility in bipolar cells, and even less in other cell types (Fig. 3B, red box). While pan BCs and ON-BCs showed nearly identical open chromatin profiles, OFF-BC open chromatin patterns were somewhat divergent, with slightly more accessibility in the photoreceptor-enriched subset (red box) and less in the bipolar-enriched subset (green box) compared to ON-BC. Overall, the sum of photoreceptor and bipolar cell ATAC-seq peaks accounted for 83 percent of whole-retina DNAse-seq peaks, with the remainder presumably deriving from other inner retinal cell classes. These data suggest that relatively few regions of open chromatin are truly photoreceptor-specific, and that regions enriched in bipolar cells are more likely to share accessibility in other tissues, such as brain. For instance, compared to photoreceptors, a greater proportion of bipolar cell peaks corresponded to DNAse-seq peaks in brain (47% vs 37%). Finally, direct comparison of ATAC-seq peaks from photoreceptor and bipolar cells identified 55,402 differentially accessible regions (FDR < 0.05), 75% of which are more accessible in bipolar cells (Table S7).

While we observed significant differences in global chromatin accessibility between photoreceptors and bipolar cells, the open chromatin profiles of ON and OFF bipolar cells were largely similar, consistent with the high degree of similarity between their transcriptomes. Specifically, only 4,263 peaks were differentially accessible between ON and OFF bipolar cells. Of note, 79% (3359) of these differential peaks were more accessible in ON bipolar cells. When examining loci surrounding genes expressed in both ON and OFF bipolar cells, we found that ON and OFF subclasses typically had similar open chromatin profiles (e.g., *Vsx2*, Fig. 3C). In contrast, some ON- or OFF-specific genes exhibit cell subclass-specific patterns of chromatin accessibility (e.g., *Isl1* and *Esam*, Fig. 3D-E). However, a correlation between gene expression and chromatin accessibility is not found at all gene loci (e.g., *Grik1*, Fig. 3F). Thus, the relationship between ON- and OFF-specific gene expression and chromatin accessibility is complex, consistent with previous observations in rods and cones (Fig S5C).

### Photoreceptor and bipolar cells employ closely related yet distinct cis-regulatory grammars

Having compared global accessibility between photoreceptor and bipolar cells, we next sought to compare them in terms of ‘*cis*-regulatory grammar’, which we define as the number, affinity, spacing and orientation of TF binding sites within the open chromatin regions of a given cell type or class. To begin, we assessed all 319 TF binding site motifs from the HOMER database^47^ for enrichment within bipolar cell open chromatin regions. The most highly enriched motifs within enhancers corresponded to CTCF, K50 HD, E-box, nuclear receptor, and MADS box motifs (Table S8). All of these motifs were previously shown to be among the most enriched motifs in photoreceptor ATAC-seq peaks as well^30^. The similarity in the patterns of TF binding site enrichment between photoreceptors and bipolar cells can be better understood in the context of known patterns of TF expression in these cell classes. Specifically, the K50 HD TFs OTX2 and CRX are master regulators of gene expression programs in photoreceptor and bipolar cells. Likewise, bHLH TFs (which recognize E-box motifs) play roles in fate specification and maintenance of both photoreceptor and bipolar cell gene expression programs, as outlined in the Introduction. Finally, enrichment of ZF motifs recognized by CTCF in both cell classes is in line with reports that CTCF motifs lie in ubiquitously accessible chromatin regions^48^, where CTCF recruitment is thought to play an architectural role, mediating the interaction between promoters and enhancers, among other functions^49,50^.

Despite the overall similarity between the *cis*-regulatory grammars of bipolar cells and photoreceptors, there are notable quantitative differences in motif enrichment between the two cell classes. To systematically identify these differences, we compared the proportion of peaks containing each of the 319 motifs between pan-bipolar cells and each photoreceptor cell type (Table S8). The most differentially enriched motifs corresponded to those with the highest enrichment in each cell class and are summarized in Figure 4. Although both photoreceptors and bipolar cells showed marked enrichment for K50 HD motifs, these motifs were more enriched in photoreceptors (Fig. 4). Conversely, E-box motifs were more enriched in bipolar cells than photoreceptors. The most striking difference in TF binding site enrichment between the two cell classes was the enrichment of both monomeric and dimeric Q50 HD motifs in photoreceptor open chromatin regions and their lack of enrichment in bipolar regions (Fig. 4). The most well-characterized Q50 HD TF expressed in photoreceptors is RAX, which is required for cone gene expression and survival^32^. In contrast, bipolar cells express multiple Q50 HD TFs (VSX2, VSX1, ISL1, LXH3, LHX4, AND SEBOX)^44^. VSX2 is required for bipolar cell development and is expressed in all mouse bipolar cell types throughout development and into adulthood^20,21^. The paradoxical absence of Q50 HD motif enrichment in bipolar open chromatin regions despite the presence of multiple Q50 HD TFs in this cell class may be explained by the observation that VSX2 acts as a repressor of photoreceptor CREs^34^. We hypothesize that the lack of enrichment of Q50 motifs in bipolar cells is due to selective closure of photoreceptor-specific open chromatin regions by VSX2, which, in turn, prevents ectopic expression of photoreceptor genes in bipolar cells. We will return to this hypothesis in the final section of the Results.

**Figure 4.**
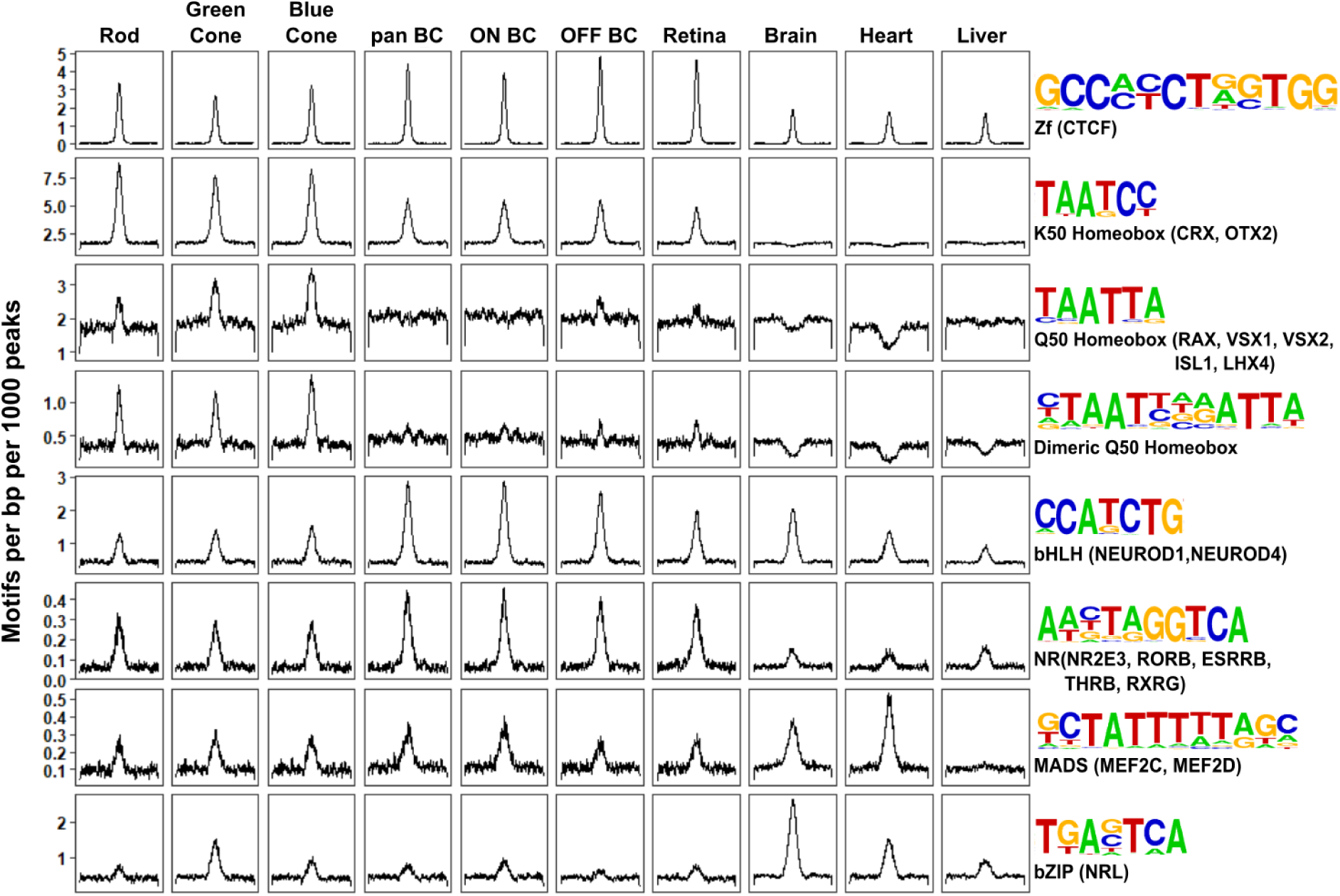
Patterns of TF binding site enrichment across retinal cell classes. Motif enrichment patterns identified in bipolar and photoreceptor ATAC-seq datasets as well as in DNAse-seq datasets from adult mouse whole retina, brain, heart and liver. This analysis included all enhancer (i.e., TSS-distal) open chromatin peaks from each cell class or tissue. Each panel is centered on a 1 kb window around peaks from the indicated dataset. Motif density (motifs per base pair per 1000 peaks) is shown on the Y-axis. Consensus sequences for each motif class and example TFs (in parentheses) expressed in photoreceptors and bipolar cells are shown on the right. Photoreceptor and bipolar cell populations share enrichment for K50 HD motifs, while only photoreceptors show enrichment for Q50 HD motifs.

To further compare the *cis*-regulatory grammars of bipolar cells and photoreceptors, we examined TF binding site co-occurrence and spacing within each cell class. For this analysis we compared a combined list of enhancer regions from rod and cones to that of bipolar cells. As previously reported for photoreceptors, motifs enriched for co-occurrence (pairs of motifs within a single open chromatin region) tended to include those that showed the highest individual enrichment^30^. For example, a common motif pair, found in open chromatin regions of both photoreceptors and bipolar cells, was a K50 HD motif co-occurring with an E-box (i.e., bHLH binding site). Overall, we found that the patterns of TF binding site co-occurrence were largely indistinguishable between bipolar cells and photoreceptors (Fig. S3)^30^. Similarly, motif co-occurrence is indistinguishable between ON and OFF bipolar cells (Fig. S3). To investigate preferences in spacing and orientation between pairs of motifs, we plotted the density of highly enriched motifs (those depicted in Fig. 4) centered on regions flanking K50 and Q50 HD motifs in each peak set. As described previously for photoreceptors, spacing and orientation preferences in bipolar open chromatin regions were minimal (Fig. S4)^30^. Thus, patterns of motif co-occurrence and spacing were largely indistinguishable between photoreceptors and bipolar cells; and the primary differences in the *cis*-regulatory grammar of the two cell classes appears to be the degree of HD and E-box motif enrichment.

### Photoreceptor- and bipolar-specific open chromatin regions are positively correlated with cell class-specific gene expression

We next sought to determine the extent to which photoreceptor- and bipolar-enriched open chromatin regions correlate with cell type-specific gene expression. To this end, we assigned each of the 55,402 regions identified as differentially accessible between photoreceptor and bipolar cells to a candidate target gene based on proximity to the nearest transcription start site and compared mean RNA-seq expression values for the assigned genes. As described in previous studies, we observed a modest correlation between enhancer accessibility and gene expression, and a more robust correlation between promoter accessibility and gene expression (Fig. S5)^30,51,52^.

Our analysis of global chromatin accessibility suggested that many of the differentially accessible peaks were also open in other tissues, especially those enriched in bipolar cells compared to photoreceptors (Fig. 3E). Therefore, to gain a better understanding of the cell type-specific open chromatin regions that drive gene expression differences between these two cell classes, we refined our analysis to exclude peaks shared with non-retinal cell types. We identified 8,435 enhancer regions which are accessible either in photoreceptors or bipolar cells, but not accessible in brain, liver or splenic B cells (Fig 5A). This set includes 1,291 regions that are open in both photoreceptors and bipolar cells, and 7,144 regions that are differentially accessible between the two cell classes (Figure 5A). We found that ~46% (3,270) of the differentially accessible peaks were more open in bipolar cells. Thus, most of the bipolar cell-enriched regions identified in the previous section were also accessible in one or more non-retinal tissues. As with the enhancer regions from the unfiltered list, assigning genes to this more retina-specific set of differentially accessible regions also shows a correlation between accessibility and gene expression (Fig. 5B). To visualize this association and identify the peaks that underly it, we plotted all 8,435 peaks according to fold-change differences in accessibility and gene expression between bipolar cells and rods (Fig. 5C) and between bipolar cells and blue cones (Fig. S5D). We then selected for further analysis those peaks that exhibited correlated accessibility and gene expression in photoreceptors (highlighted in red or blue in Fig. 5C and S5D; n = 901) or bipolar cells (highlighted in green in Fig. 5C and S5D; n = 833). These differentially enriched peaks represent strong candidates for CREs that mediate the gene expression differences between the two cell classes. Indeed, the photoreceptor peak set contains known enhancers responsible for driving cell type-specific expression of *Rhodopsin* (*Rho*) and components of the rod-specific phototransduction cascade^19,53^, while the bipolar peak set contains a known enhancer driving *Vsx2* expression in bipolar cells^54^. To gain insight into the possible biological functions of these peaks we used GREAT^55^ to assign biological annotations based on nearby genes and identified highly enriched biological processes associated with ‘sensory perception of light stimulus’ in the photoreceptor peak set and ‘transmission of nerve impulse’ in the bipolar peak set (Fig. S6). Thus, as was found with the unfiltered datasets, photoreceptor peaks are linked with genes associated with light sensation, whereas bipolar peaks are linked to genes involved in more generic neuronal functions.

**Figure 5.**
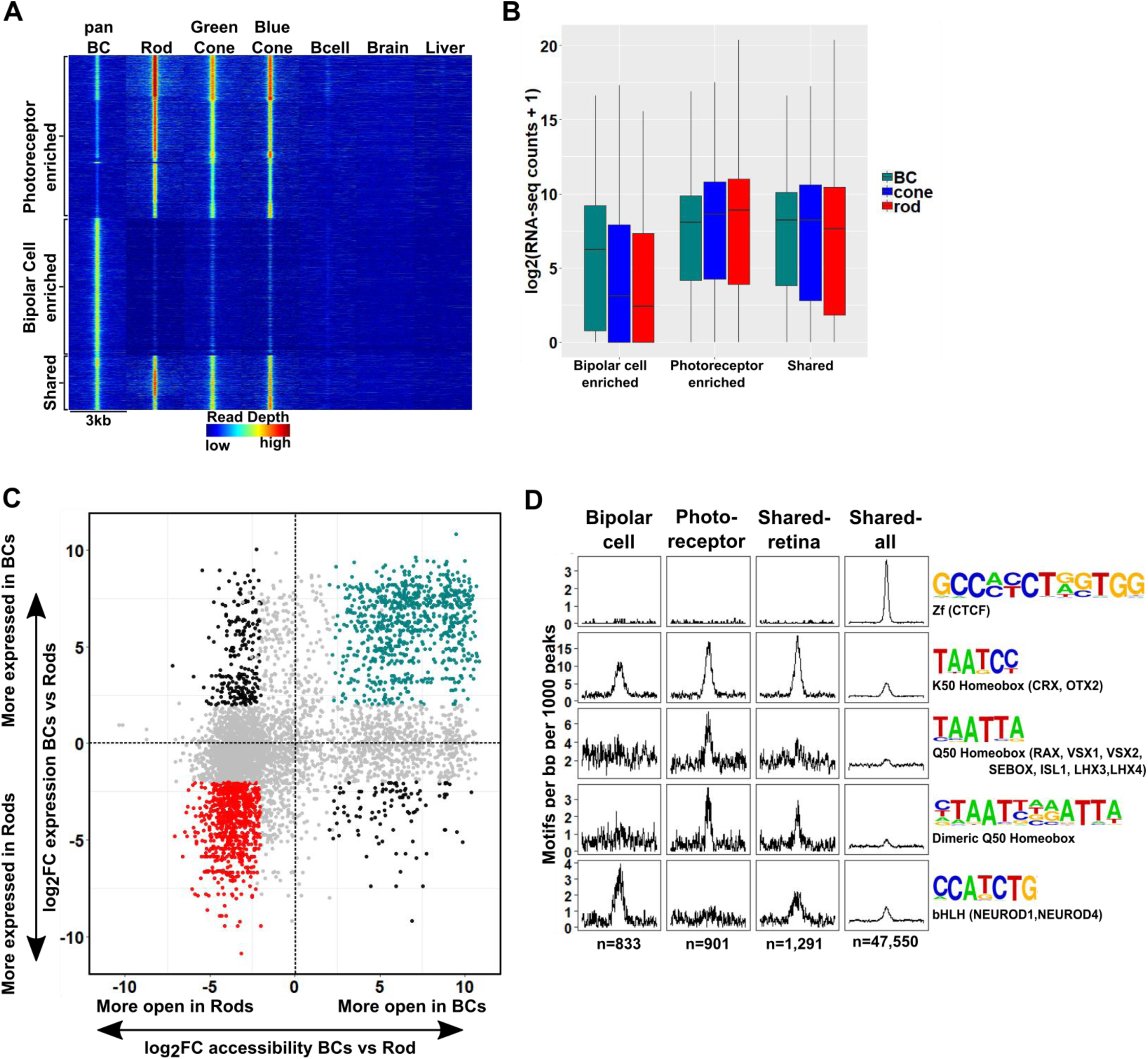
The *cis*-regulatory grammar of genomic regions associated with differential chromatin accessibility and gene expression in bipolar cells and photoreceptors. (**A**) Heatmap depicting 8,435 genomic regions determined by pairwise comparison to be differentially accessible in photoreceptors, bipolar cells, or both cell classes, compared to adult mouse B cells, brain and liver. Photoreceptor-enriched (n = 3,874), bipolar cell-enriched (n = 3,270), shared (n = 1,291). (**B**) Expression of genes to which classes of peaks defined in A were assigned by proximity to nearest TSS. There is a moderate correlation between chromatin accessibility and gene expression in each cell class. (**C**) Peaks identified in panel A are plotted according to chromatin accessibility (x-axis) and associated gene expression (y-axis) in bipolar cells versus rods. Peaks with four-fold greater chromatin accessibility and associated gene expression in bipolar cells are shown in green (FDR < 0.05 for both accessibility and expression), while those peaks with greater accessibility and associated gene expression in rods are shown in red. Shared peaks and those associated with modest (less than four-fold) differences in expression are in grey. Peaks with discordant chromatin accessibility and associated gene expression are shown in black. (**D**) Motif enrichment within peaks displaying correlated chromatin accessibility and associated gene expression in bipolar cells and photoreceptors as well as within peaks displaying shared accessibility, including those shown in panel A (shared-retina) and shared peaks which were not filtered to remove those accessible in non-retinal tissues (shared-all). Both photoreceptor and bipolar cell peaks show enrichment for K50 HD motifs. Bipolar cell peaks (n = 833) are also highly enriched for E-box motifs but lack enrichment of Q50 HD motifs. In contrast, photoreceptor peaks (n = 901) show enrichment for Q50 HD motifs but almost entirely lack E-box enrichment. Shared retina-specific peaks (n = 1,291) show a hybrid pattern of motif enrichment. Only the shared-all peak set exhibits enrichment for CTCF motifs, underscoring a key difference between cell-class specific open chromatin regions (which show no enrichment of CTCF motif) and ubiquitously open chromatin regions which show strong enrichment.

Next, we asked whether the patterns of TF binding site enrichment observed with aggregate sets of ATAC-seq peaks from each cell class would be preserved within the retina-specific peak sets associated with correlated gene expression. We compared photoreceptor-enriched regions, bipolar-enriched regions, and regions that share accessibility between the two cell classes which were either specific to the retina (Fig. 5A, ‘shared retina’ n = 1,291), or unfiltered (‘shared all’, n = 47,550). We found that K50 HD motifs were enriched in both shared and cell class-selective regions, but to a lesser extent in regions specifically enriched in bipolar cells. The bipolar-selective regions were markedly enriched for E-box motifs but completely lacked enrichment for Q50 HD motifs (Fig. 5D). Conversely, photoreceptor-selective regions were enriched for Q50 HD motifs, but lacked E-box motif enrichment. Peaks that were shared between the two cell classes showed an intermediate pattern of motif enrichment. Of note, CTCF enrichment was absent in all but the unfiltered peak set, suggesting that the CTCF enrichment observed in Figure 4 is attributable to ubiquitously accessible peaks. Taken together, these findings suggest that differential enrichment of Q50 HD and E-box motifs are the key features that distinguish the *cis*-regulatory grammars of photoreceptors and bipolar cells.

### K50 motifs in the Gnb3 promoter are required for both photoreceptor and bipolar expression, but addition of Q50 motifs selectively represses expression in bipolar cells

Given the critical roles of HD TFs in the regulation of photoreceptor and bipolar gene expression, we further investigated the role of K50 and Q50 motifs in a specific *cis*-regulatory region, the promoter of *Gnb3*. *Gnb3* encodes the β subunit of a heterotrimeric G-protein required for cone phototransduction as well as ON bipolar cell function. *Gnb3* is expressed in rods, cones, and bipolar cells during early postnatal retinal development in the mouse. Selective repression of *Gnb3* in rods by the nuclear receptor TF NR2E3 results in a cone + bipolar pattern after postnatal day 10^56,57^. We focused on the 820 bp immediately upstream of the TSS of *Gnb3* which drives robust expression in rods, cones, and bipolar cells when electroporated into early postnatal mouse retina. This region lacks Q50 motifs but contains five K50 HD motifs of varying affinity which occur in two clusters, one immediately upstream of the TSS (−65 bp) and the other more distally (−350 bp). To evaluate the role of these five K50 motifs in mediating photoreceptor and bipolar expression, we engineered reporter constructs in which each of the five motifs was individually inactivated by mutating the TAAT core to TGGT. We then introduced wild-type and mutant reporters into mouse retinal explants via electroporation and compared expression levels after 8 days. Mutations in K50 motifs 2, 4 and 5 resulted in coordinate loss of expression in both photoreceptor and bipolar cells, indicating that these motifs are required for reporter expression in both cell classes. Conversely, mutations in site 1 or 3 had no effect on expression in either cell class (Fig. 6B). Binding site affinity did not correlate with expression, as site 3 has a higher predicted affinity than sites 4 or 5. Thus, the *Gnb3* promoter contains both essential and nonessential K50 motifs, underscoring the critical role for these shared motifs in both photoreceptors and bipolar cells.

**Figure 6.**
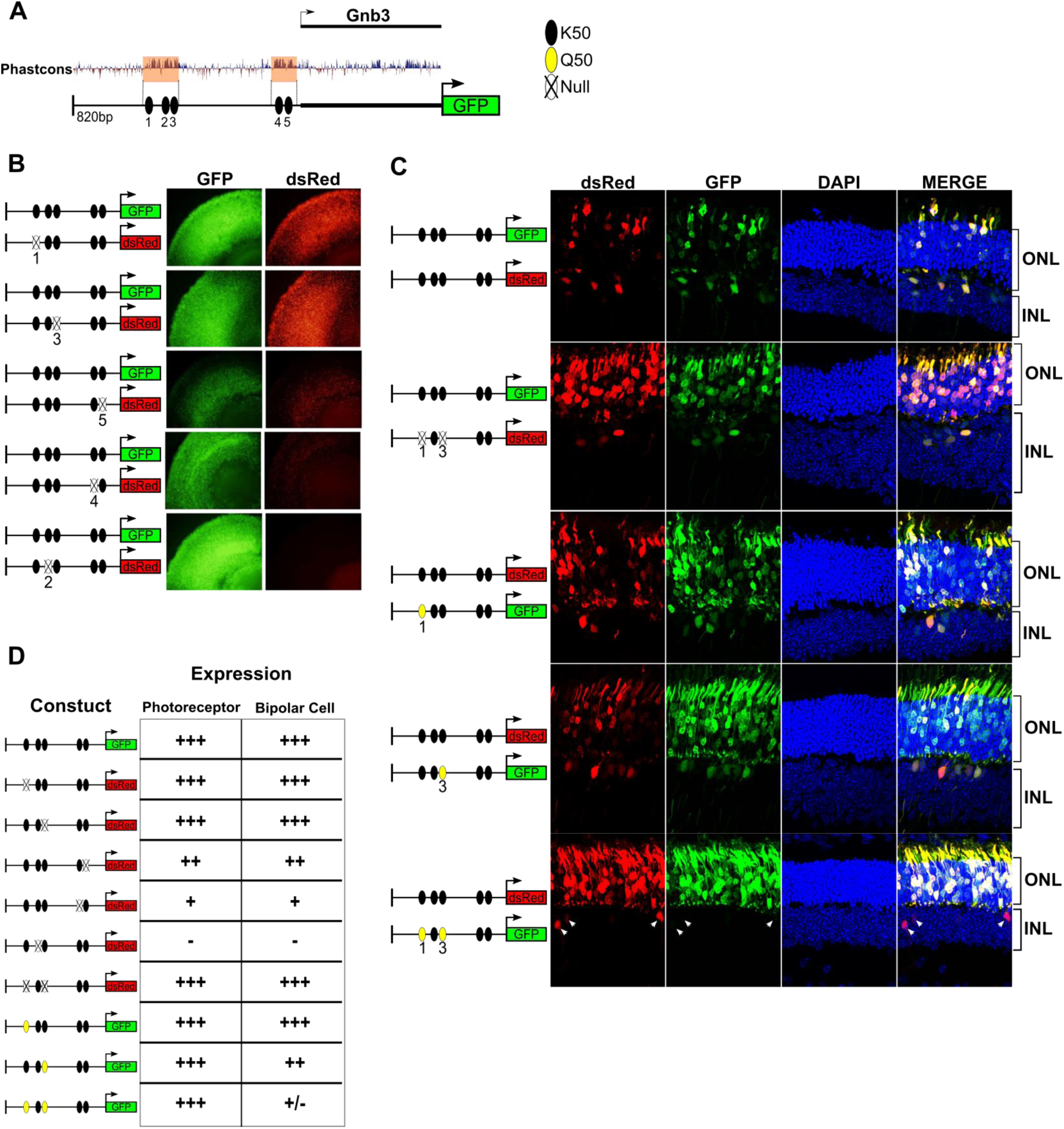
K50 motifs are required for expression in both photoreceptors and bipolar cells, while Q50 motifs mediate repression of reporter expression specifically in bipolar cells. (**A**) Schematic of reporter construct containing 820 bp from the promoter region and 5’ untranslated region (UTR) of mouse *Gnb3*. This region contains two phylogenetically conserved blocks harboring a total of five K50 motifs (black ellipses). (**B**) **Left**, Schematics of reporter pairs containing wild-type (WT; black) and inactivated (crossed out) K50 motifs. **Right**, Each pair of reporters was electroporated into explanted newborn mouse retina, which were subsequently harvested after eight days in culture and photographed in a flat mount preparation. While K50 sites 2, 4 and 5 are required for reporter expression, sites 1 and 3 are dispensable. (**C**) **Left**, Pairs of reporters containing WT, inactivated K50, or novel Q50 (TAATTA; yellow) motifs were electroporated into newborn mouse retina *in vivo*. Mice were allowed to grow for 20 days at which point retinas were harvested and photographed in vertical cross-sections. **Right**, Representative cross-sections of retinas injected with the indicated pair of reporters. Loss of both K50 motifs 1 and 3 has no effect on expression in either photoreceptors or bipolar cells, while converting these same sites to Q50 motifs abrogates expression specifically in bipolar cells (n = 3-6 depending on reporter pair). White arrowheads indicate bipolar cells expressing the WT reporter, but not the mutated one. (**D**) Table summarizing the results of reporter analysis presented in (**B**) and (**C**).

Next, we sought to determine the effect of introducing Q50 motifs into the *Gnb3* promoter. For these experiments, reporters were introduced into newborn mouse retinas via *in vivo* electroporation and harvested for histologic analysis after 20 days. First, we electroporated identical wild-type sequences driving both dsRed and GFP to confirm that essentially all photoreceptors and bipolar cells received both constructs (Fig. 6C). Next, we compared the expression of a *Gnb3* promoter containing mutations in K50 motifs 1 and 3 to that of a wild-type promoter, confirming that elimination of both of these sites has no effect on expression in either photoreceptors or bipolar cells (Fig. 6 C, D). To test the effect of introducing Q50 motifs into the *Gnb3* promoter, we replaced K50 motifs 1 and 3 with a Q50 motif (TAATTA), both individually and in combination. Whereas introduction of a Q50 motif into site 1 had no apparent effect, replacement of site 3 caused a selective decrease in bipolar expression with no change in photoreceptor expression. When we introduced Q50 motifs into both sites, reporter expression was selectively turned off in bipolar cells, with no effect on photoreceptor expression. These data, along with previous reports of VSX2-mediated repression of photoreceptor-specific promoters/enhancers, suggest that Q50 motifs play an important role in mediating repression of photoreceptor genes in bipolar cells.

## Discussion

In this study we generated open chromatin maps and transcriptome profiles of mouse bipolar cells, including FACS-purified ON and OFF bipolar cell populations, and compared them to analogous data from rod and cone photoreceptors. We found that photoreceptors and bipolar cells differ in the expression of thousands of genes, and yet have very similar *cis*-regulatory grammars. The key *cis*-regulatory differences that distinguish the two cell classes are the preferential enrichment of Q50 HD motifs in open chromatin regions associated with photoreceptor-specific gene expression and a corresponding enrichment of E-box motifs in chromatin associated with bipolar-specific expression. The cellular features and transcriptional mechanisms shared by photoreceptors and bipolar cells have prompted speculation that these two sister cell types arose from a single ancestral photoreceptor cell type via a process of progressive cellular divergence^5,6^. We propose that the elimination of Q50 motifs from bipolar-specific CREs likely played a key role in differentiating the bipolar transcriptome from that of photoreceptors during early stages of vertebrate retinal evolution. Prior studies of individual photoreceptor CREs showed a role for the Q50 HD TF, VSX2, in selectively repressing photoreceptor genes in bipolar cells^20,34^. Our results generalize this conclusion, suggesting that VSX2 plays a genome-wide role in silencing photoreceptor gene expression in bipolar cells. A similar role for VSX2 has recently been described in the spinal cord, where closely related progenitor cells give rise to either motor neurons or V2a interneurons^58^. VSX2 promotes V2a identity by directly repressing the motor neuron gene expression program and by competing for Q50 sites at motor neuron enhancers. Thus, in both retina and spinal cord, expression of VSX2 promotes interneuron fate at the expense of the alternative ‘effector’ neuron (photoreceptor or motor neuron) cell type. These parallels suggest that transcriptional repression by cell type-specific TFs such as VSX2 represent a common mechanism for differentiating the gene expression programs of two closely related cell types.

Support for the idea that bipolar cells diverged from photoreceptors via progressive partitioning of cellular function is provided by the existence of cell types in the retinas of non-mammalian vertebrates with features intermediate between those of mammalian photoreceptors and bipolar cells. In some turtle species ~30% of the cell bodies in the photoreceptor layer (the outer nuclear layer) belong to bipolar cells, not photoreceptors^12^. These so-called ‘displaced bipolar cells’ possess an inner segment-like process that extends to the outer limiting membrane and contains abundant mitochondria and even a sensory-type (‘9 + 0’) cilium (Fig. 7A,B). Thus, displaced bipolar cells closely resemble typical photoreceptors except that they lack an outer segment, possess dendrites in the outer plexiform layer, and synapse directly onto retinal ganglion cells. Another intermediate type of bipolar cell occurs in nearly all non-mammalian vertebrate classes and even in some mammalian species^13–15,59^. This bipolar type has a nucleus localized to the inner nuclear layer, but retains an inner segment-like structure, called Landolt’s club, which extends from the cell’s dendritic arbor to the outer limiting membrane and contains abundant mitochondria and a sensory-type cilium (Fig. 7A)^13–15^. We suggest that displaced bipolar cells and those with a Landolt’s club represent ‘transitional forms’ on the evolutionary path from photoreceptor to typical bipolar cell. The existence of these transitional forms suggests that bipolar cells may have evolved via the stepwise repression of discrete gene modules required for the development of individual cellular features, or ‘apomeres’, that are specific to photoreceptors.

**Figure 7.**
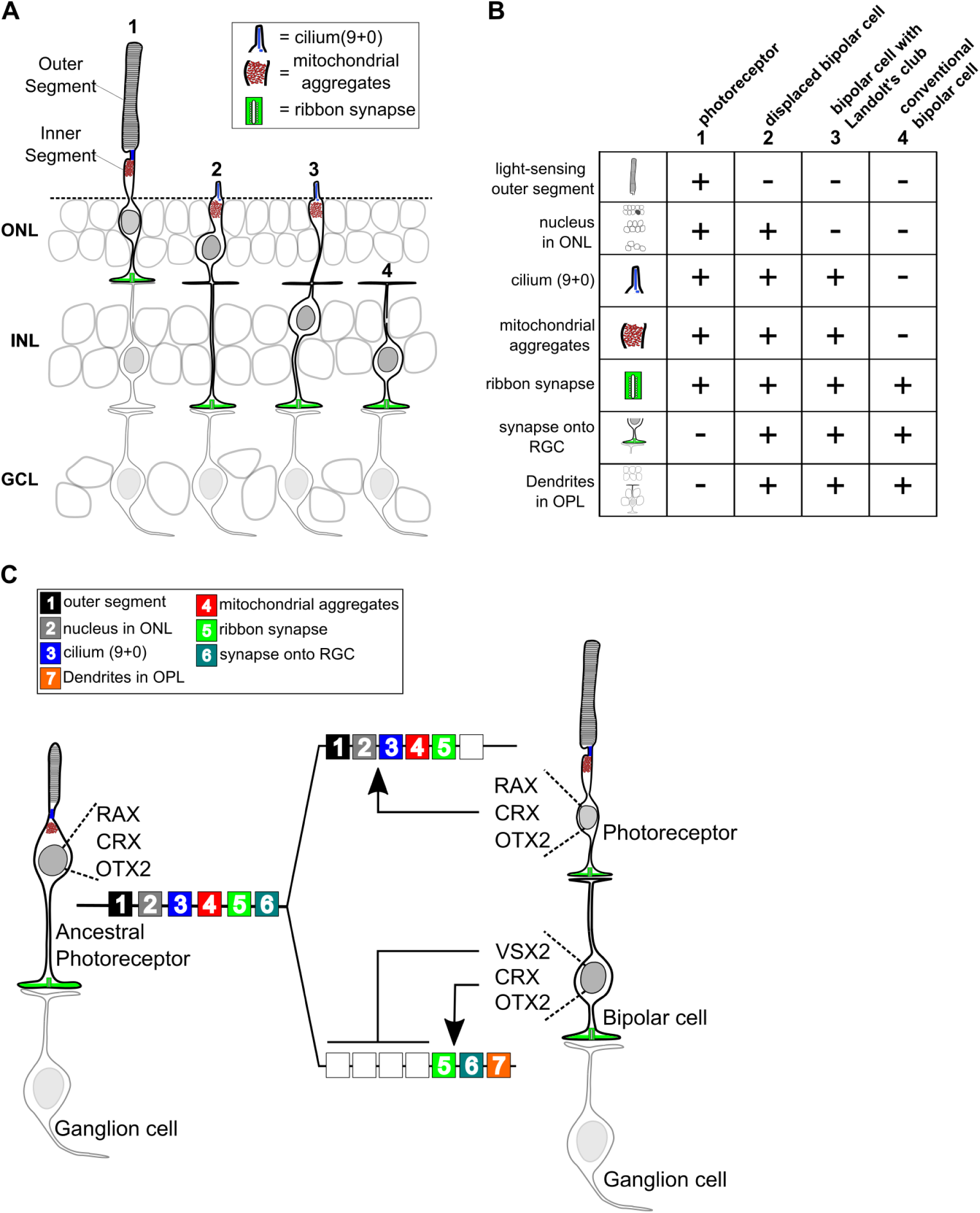
Evolutionary model for the divergence of bipolar cells from photoreceptors. (**A**) Schematic depiction of photoreceptors (1), conventional bipolar cells (4), and two ‘transitional’ cell types with cellular features intermediate between those of photoreceptors and conventional bipolar cells: displaced bipolar cells (2) and bipolar cells with Landolt’s club (3). (**B**) Table of individual cellular features (referred to in Arendt *et al.*^4^ as ‘apomeres’) possessed by photoreceptors, bipolar cells, or both cell classes. (**C**) Evolutionary model for the divergence of present-day photoreceptors and bipolar cells from a common ancestral photoreceptor type. We propose that the ancestral photoreceptor (possibly present in a hagfish-like ancestor) expressed cell type-specific genes via both K50 HD TFs (CRX and OTX2) and a possibly weakly activating Q50 HD TF (RAX). The emergent expression of a strongly repressive Q50 HD TF (VSX2) in bipolar cells then permitted silencing of selected photoreceptor gene modules underlying the formation of defined photoreceptor apomeres (e.g., outer segment). Selective expression of activating Q50 HD TFs in ‘transitional’ bipolar cell types may have allowed the derepression of specific photoreceptor apomeres (e.g., cilium formation, mitochrondrial aggregates). Novel bipolar-specific apomeres (e.g., dendrites in the outer plexiform layer [OPL]) may have evolved via co-option of other gene expression programs.

If this evolutionary model is correct, then how can we account for the co-existence of ‘transitional’ bipolar cell types and ‘conventional’ bipolar cells in a single retina? One testable hypothesis is that VSX2 may be expressed at lower levels in transitional bipolar cell types, thereby permitting expression of additional photoreceptor gene modules and their corresponding apomeres. Alternatively, it is possible that additional activating Q50 HD TFs are expressed in transitional bipolar types, and these TFs can overcome VSX2-mediated repression of selected photoreceptor gene modules. Indeed, we have found that multiple Q50 HD TFs are expressed in subsets of mouse bipolar cells. In addition, there is evidence that transitional bipolar cell types with Landolt’s club may exist in the mouse^60^. Thus, individual bipolar cell types may control the number of photoreceptor apomeres they express by modulating the balance of activating and repressing Q50 HD TFs in their nuclei.

The evolutionary divergence of bipolar cells from photoreceptors likely required coordinated changes in both *cis*-regulatory grammar and HD TF expression. The Q50 HD TF, RAX, is expressed in developing vertebrate rods and cones and is required for normal activation of photoreceptor gene expression in mice^32,61^. The expression of a RAX homolog in the photoreceptors of the tadpole larva of the protochordate, *Ciona intestinalis*, suggests a primordial role for this Q50 HD TF in activating photoreceptor gene expression in chordates. These data suggest that both K50 and Q50 motifs were present in the CREs of the ancestral vertebrate photoreceptor prior to the evolutionary emergence of bipolar cells, and that both K50 (OTX2 and CRX) and Q50 (RAX) HD TFs were required for gene activation in that ancestral cell type (Fig. 7C). In this context, the emergent expression of a repressive Q50 HD TF (VSX2) in a primordial bipolar cell would have permitted selective repression of CREs containing Q50 motifs. Maintaining expression of selected ‘photoreceptor’ genes in bipolar cells (e.g., *Gnb3*) would then have required the elimination of Q50 motifs from the *cis*-regulatory regions of those genes.

These evolutionary considerations suggest that the modern vertebrate retina arose from an ancestral retina in which photoreceptors directly synapse onto projection neurons (i.e., ganglion cells) without an intervening layer of interneurons (left side of Fig. 7C). Two lines of evidence suggest that such a retina may have existed. First, the retina of the hagfish, the most primitive extant vertebrate, reportedly has photoreceptors that directly synapse onto projection neurons (i.e., ganglion cells)^6,62^. Second, some vertebrate species (including reptiles, amphibians, and larval lamprey) have an unpaired, median ‘parietal eye’ developmentally related to the pineal gland, which contains photoreceptors that directly synapse onto ganglion cells^63–66^. It is possible that the parietal eye evolved from the midline ‘eye’ of a protochordate ancestor, akin to the present-day ascidian larva. The simple lateral eyes of a hagfish-like vertebrate ancestor may then have emerged via co-option of the gene networks required for parietal eye development. Subsequently, bipolar cells may have arisen in the lateral eyes of early vertebrates via subtle changes in *cis*-regulatory grammar and TF expression, paving the way for the emergence of the sophisticated interneuronal circuitry found in present-day vertebrate retinas.

---

**Supplementary Figure S1.**
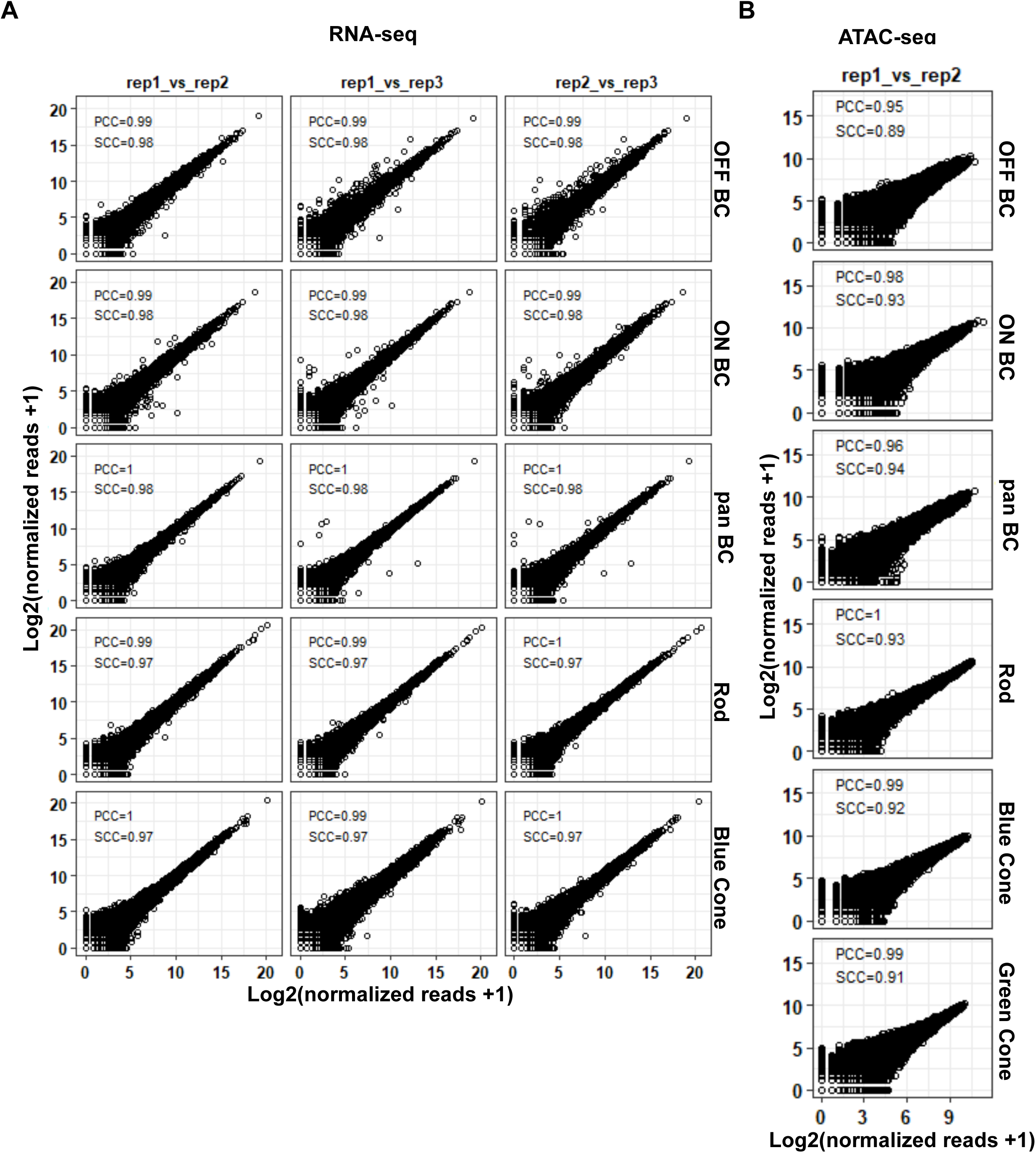
Reproducibility of ATAC-seq and RNA-seq datasets. (**A**) Pairwise comparison between biological replicates of RNA-seq from purified photoreceptor and bipolar cell populations. For each sample, normalized read counts from one biological replicate are plotted against normalized read counts of another biological replicate. (**B**) Pairwise comparison between biological replicates of ATAC-seq from purified photoreceptor and bipolar cell populations as in A. PCC: Pearson Correlation Coefficient. SCC: Spearman Correlation Coefficient.

**Supplementary Figure S2.**
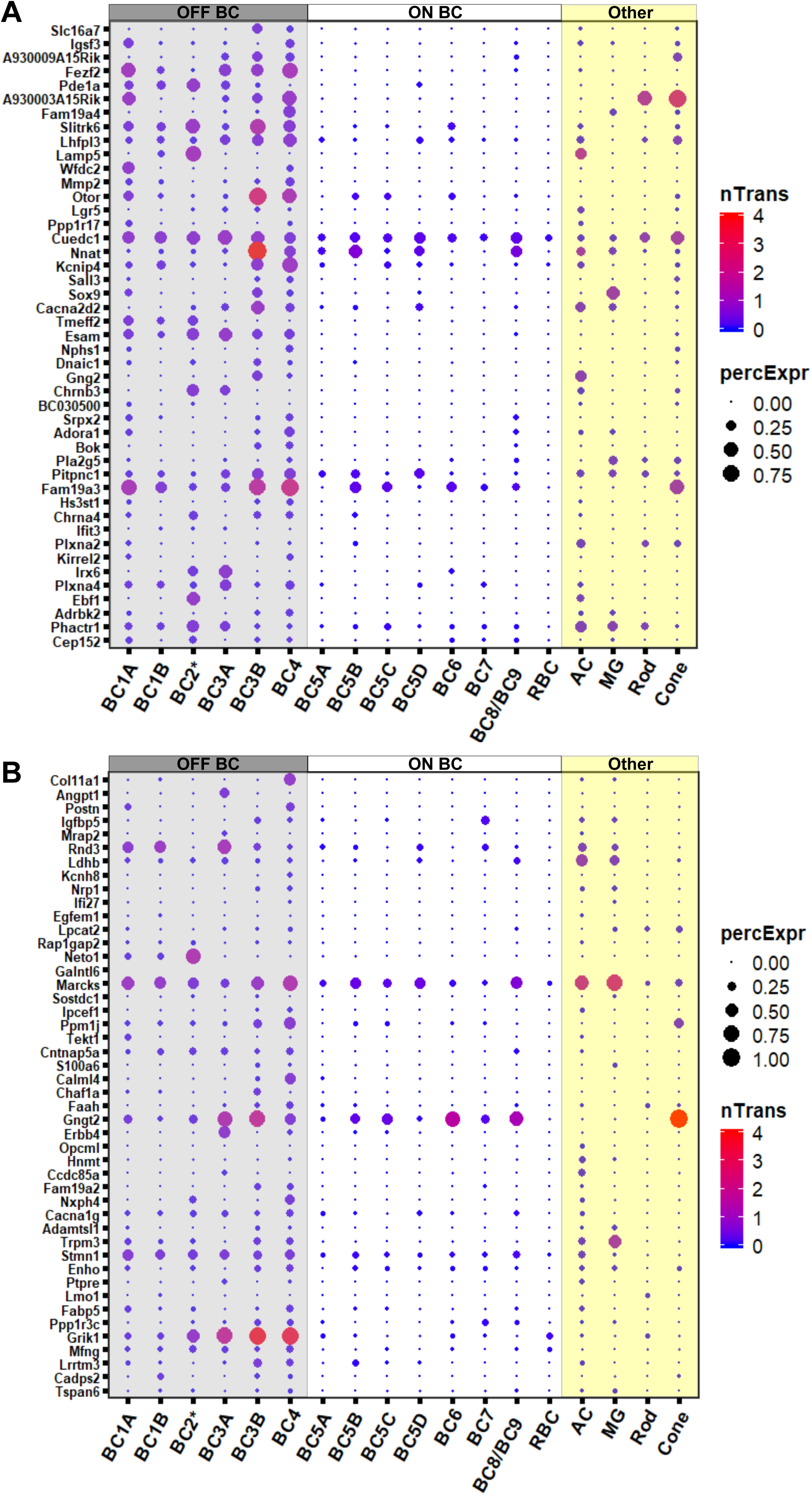
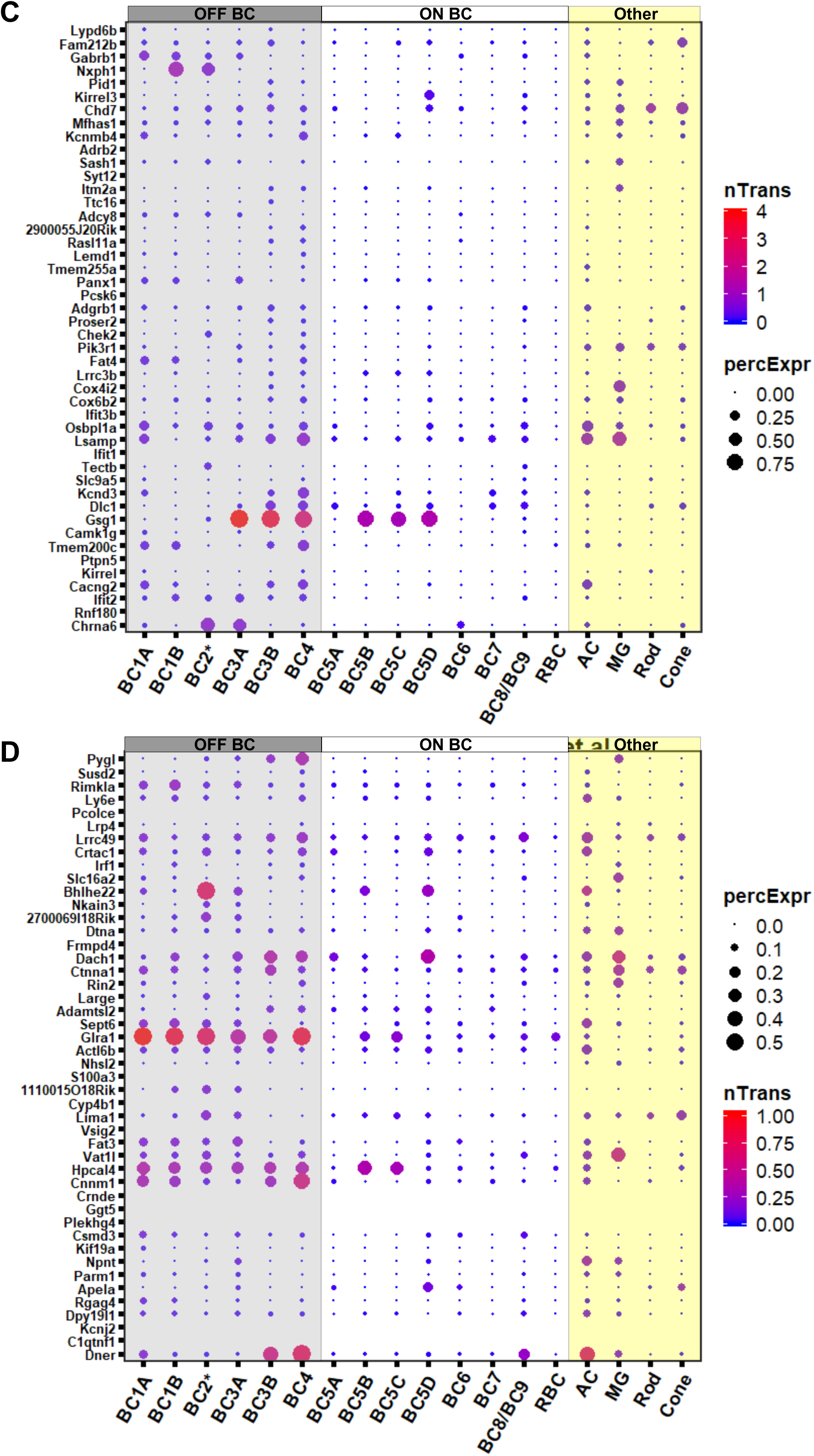
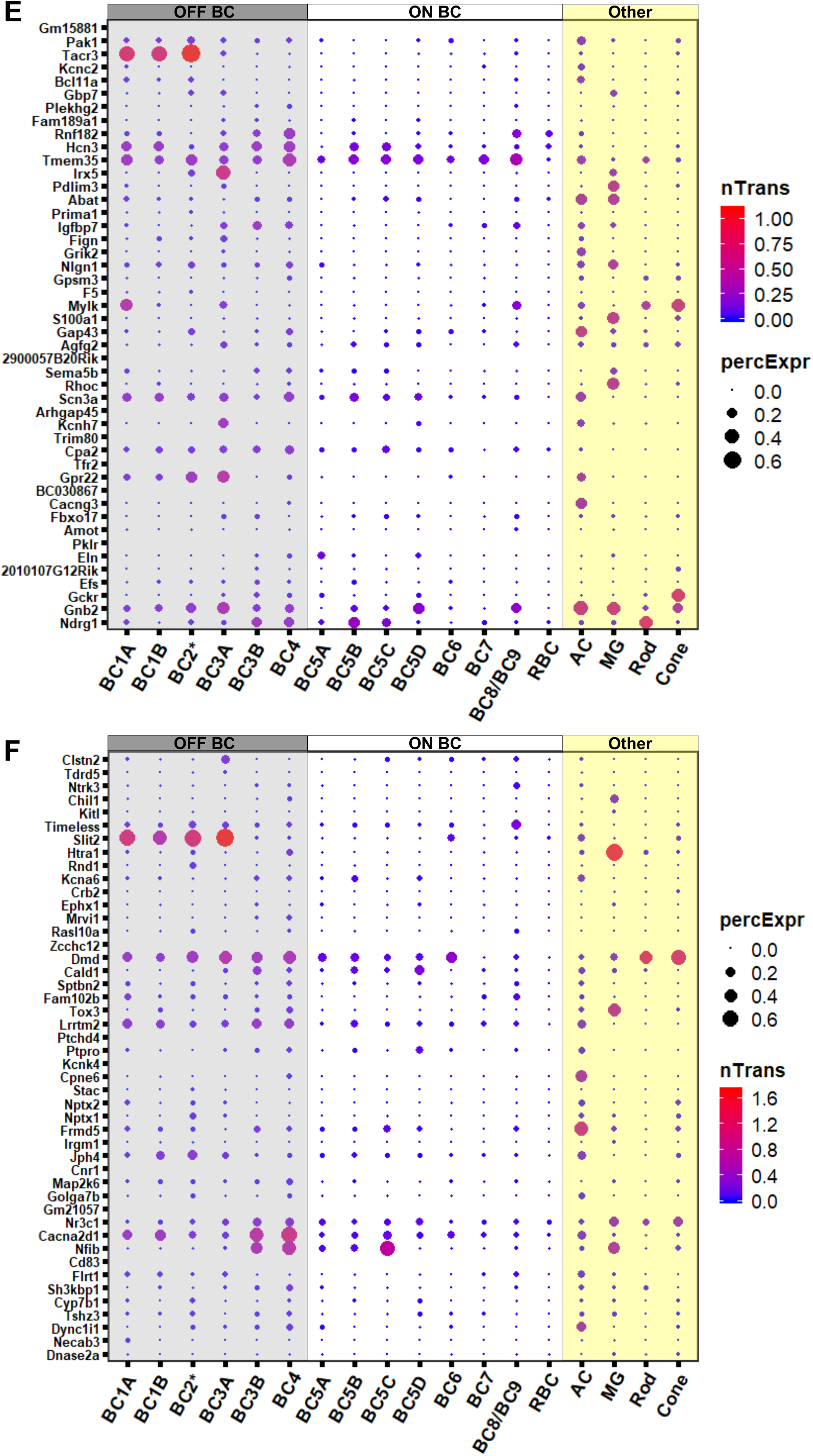
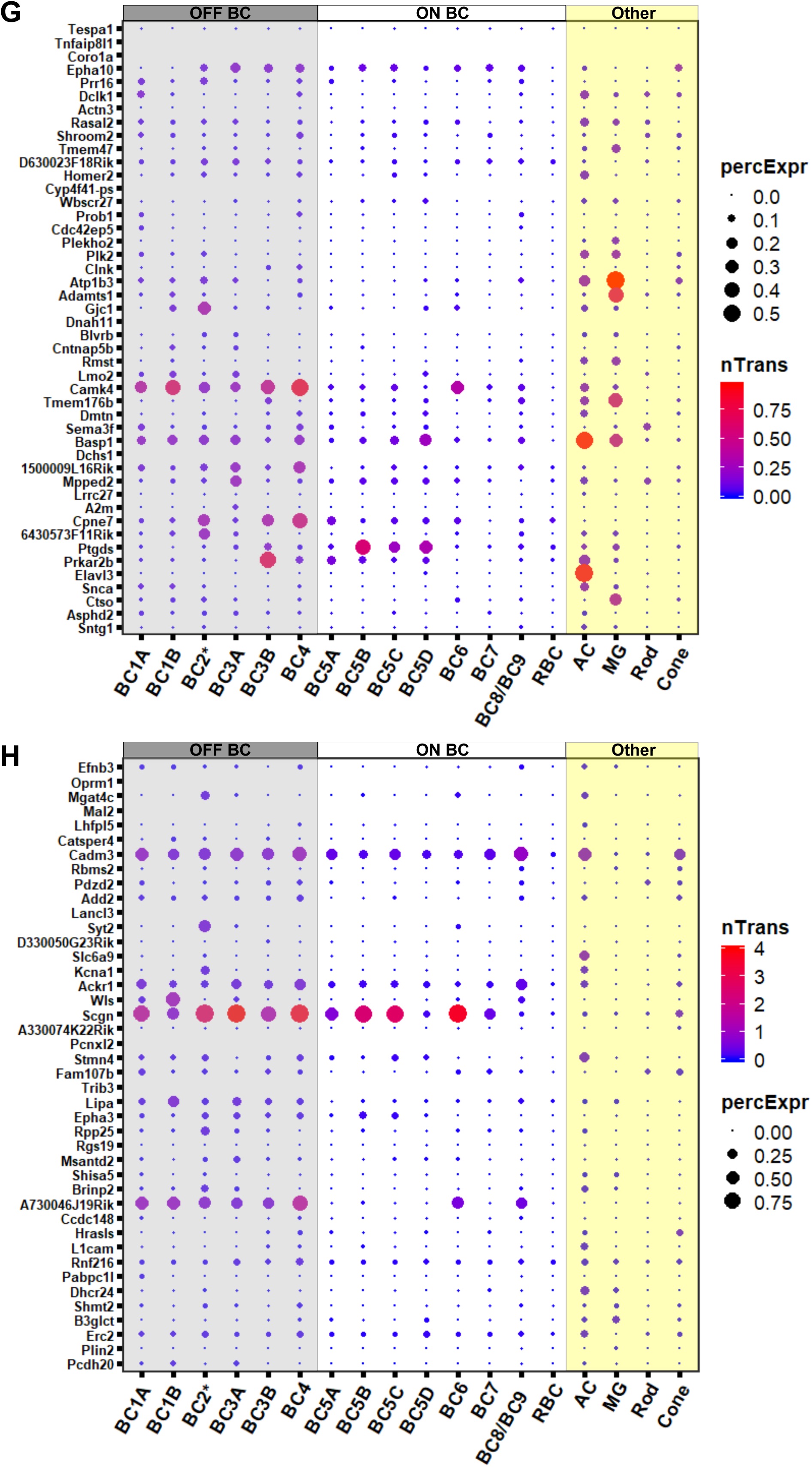
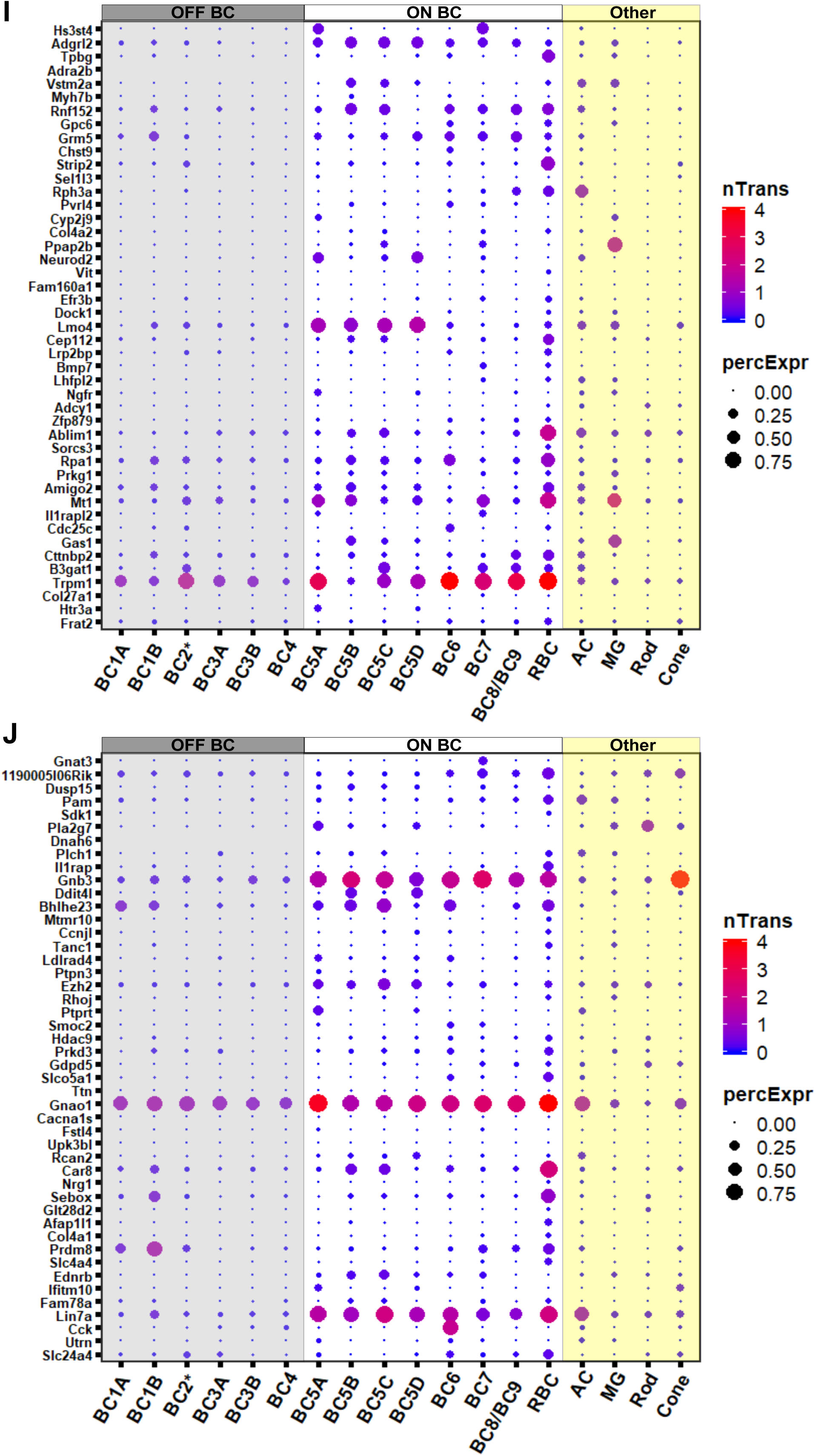
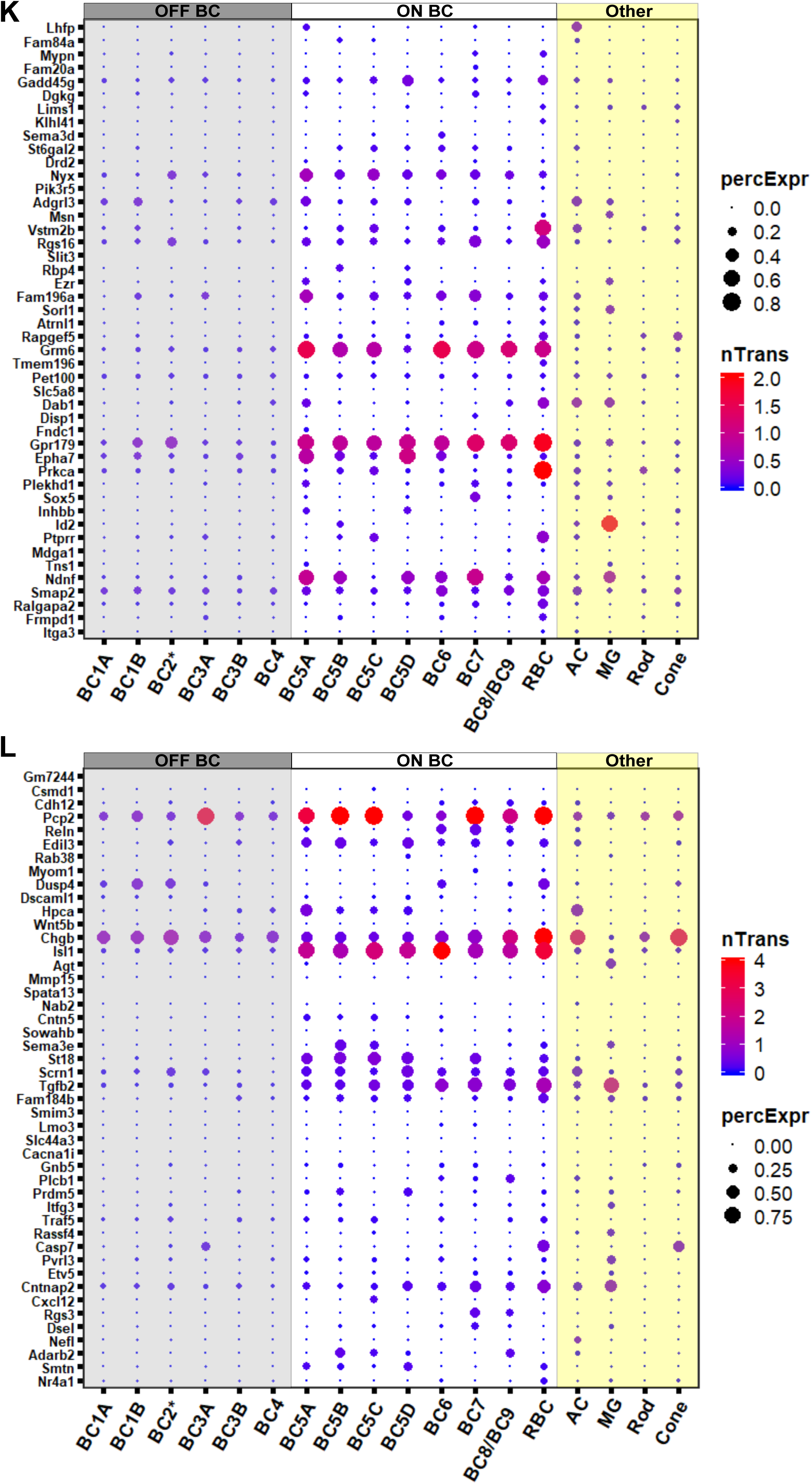
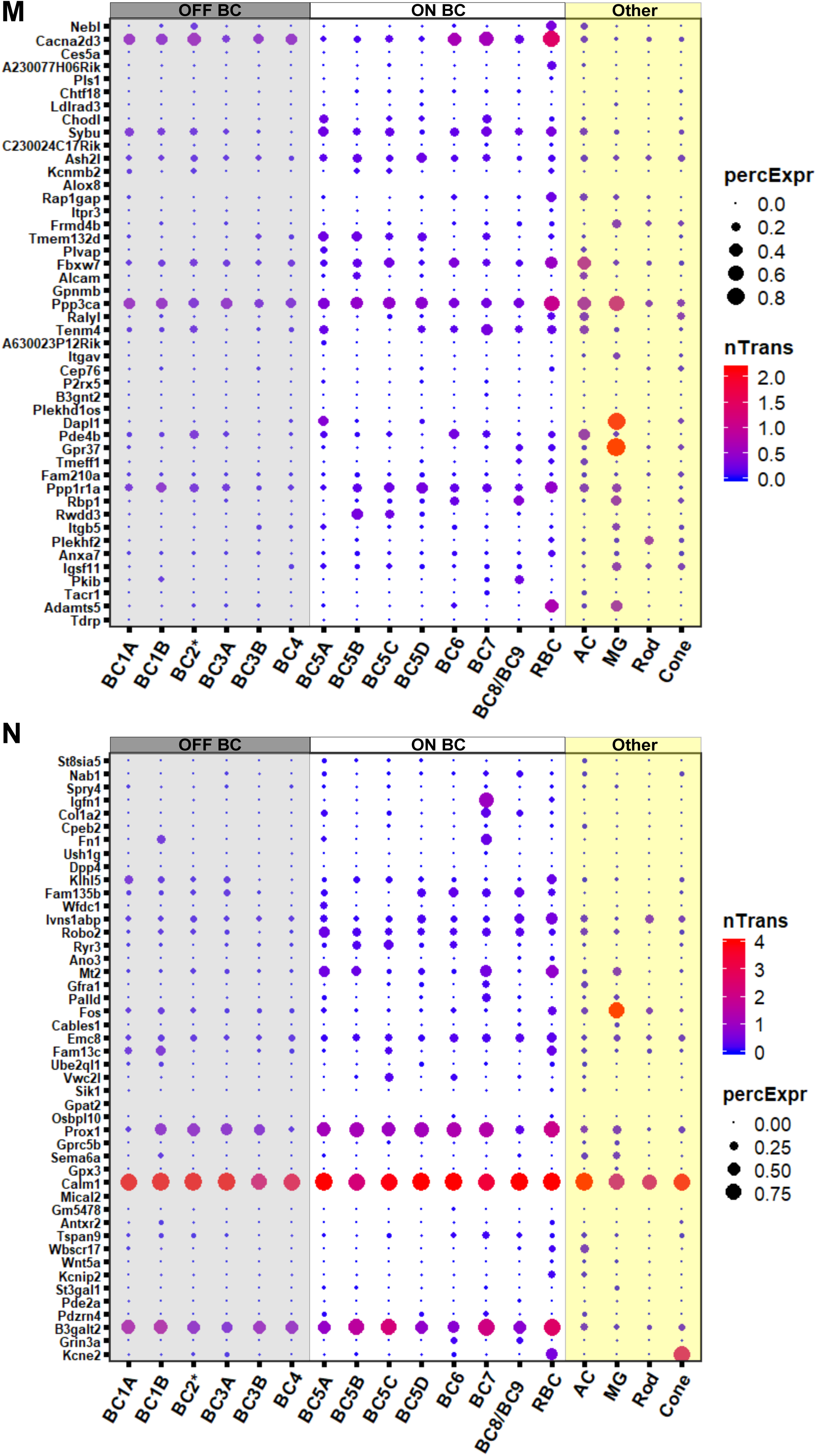
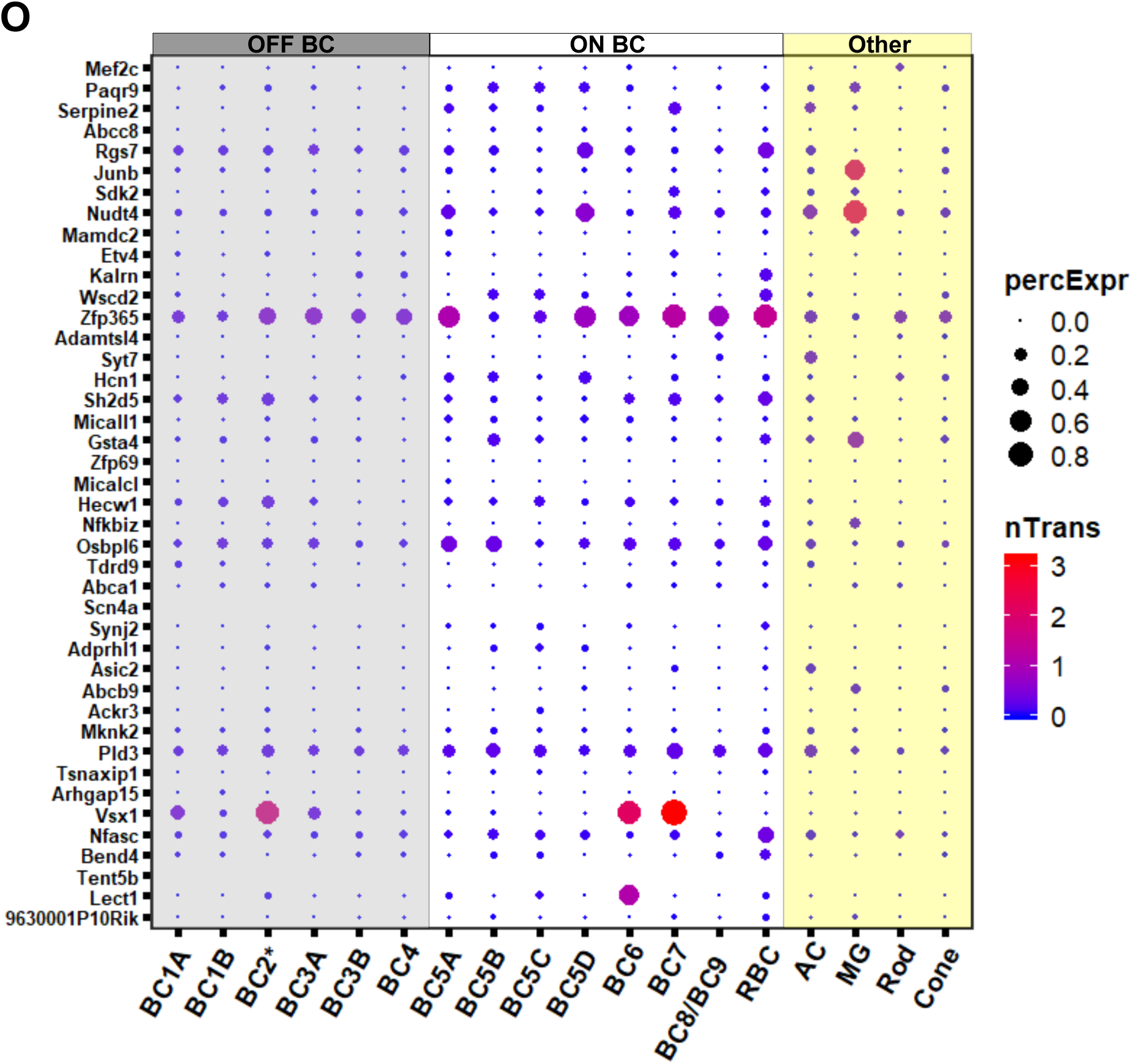
Single-cell expression profiles of genes differentially expressed between ON and OFF bipolar cells. Dot plots present single-cell expression data obtained by Shekhar *et al*.^44^ using Drop-seq for genes identified in this study by bulk RNA-seq as differentially enriched in ON or OFF bipolar cells. Drop-seq data was available for 630 of the 680 genes, which are sorted by lowest adjusted p-value. (**A-H**) Genes enriched in OFF bipolar cells. (**I-O**) Genes enriched in ON bipolar cells. nTrans = mean number of transcripts expressed per cell in each cluster identified as a bipolar cell type. PercExpr = percentage of cells within each cluster found to express the indicated gene.

**Supplementary Figure S3.**
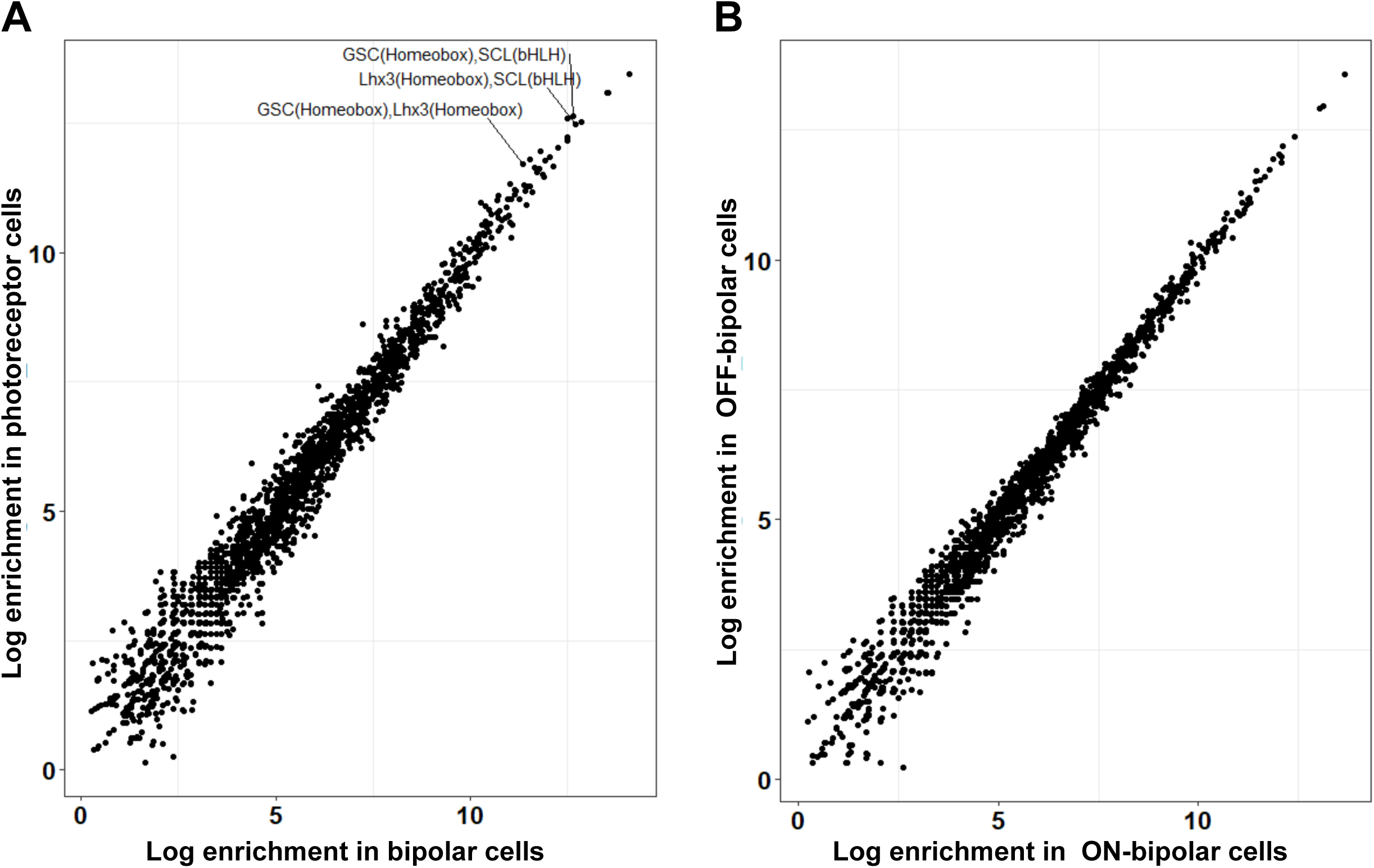
Co-enrichment of motifs in photoreceptor and bipolar cells. (**A**) Scatterplot comparing enrichment for pairs of 67 non-redundant motifs from HOMER within enhancer (TSS-distal) regions of photoreceptors (y-axis) and bipolar cell (x-axis). Each point represents a motif pair (motifs present within the same peak). Motif pairs that are highly enriched in both tissues include individual motifs recognized by GSC (K50 HD), LHX3 (Q50 HD) and SCL (E-box). (**B**) Scatterplot as in A, showing enrichment of motif pairs between ON and OFF bipolar cells.

**Supplementary Figure S4.**
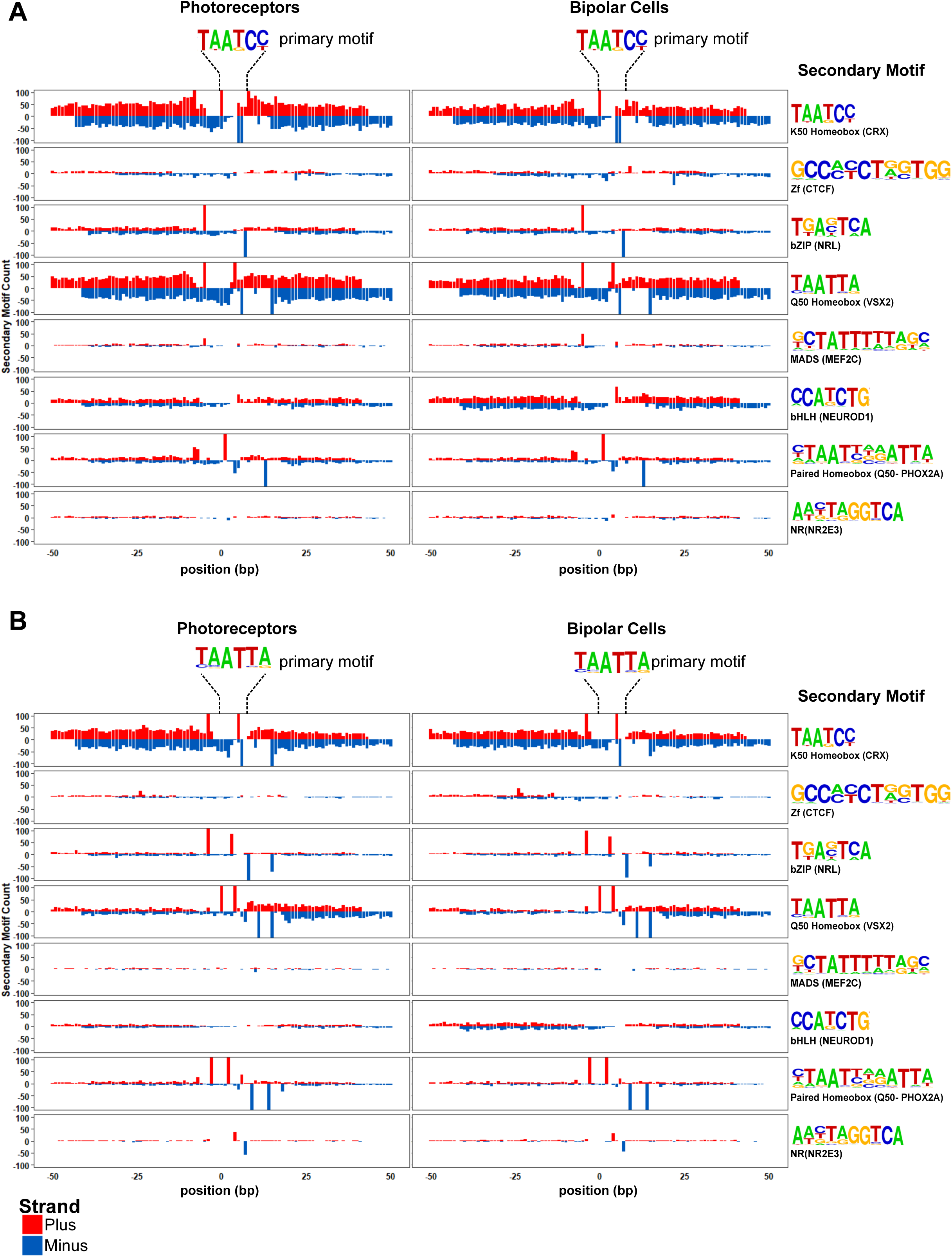
Motif spacing and orientation in photoreceptor and bipolar cells. Peaks from photoreceptors (left) and bipolar cells (right) are centered on K50 HD (**A**) or Q50 HD motifs (**B**). Per base-pair counts for secondary motifs (PWM and example TFs shown at the far right) are plotted in a 100 bp window surrounding the primary motif, with counts on the plus strand shown in red, ad the minus strand shown in blue. The y-axis was limited to ±100.

**Supplementary Figure S5.**
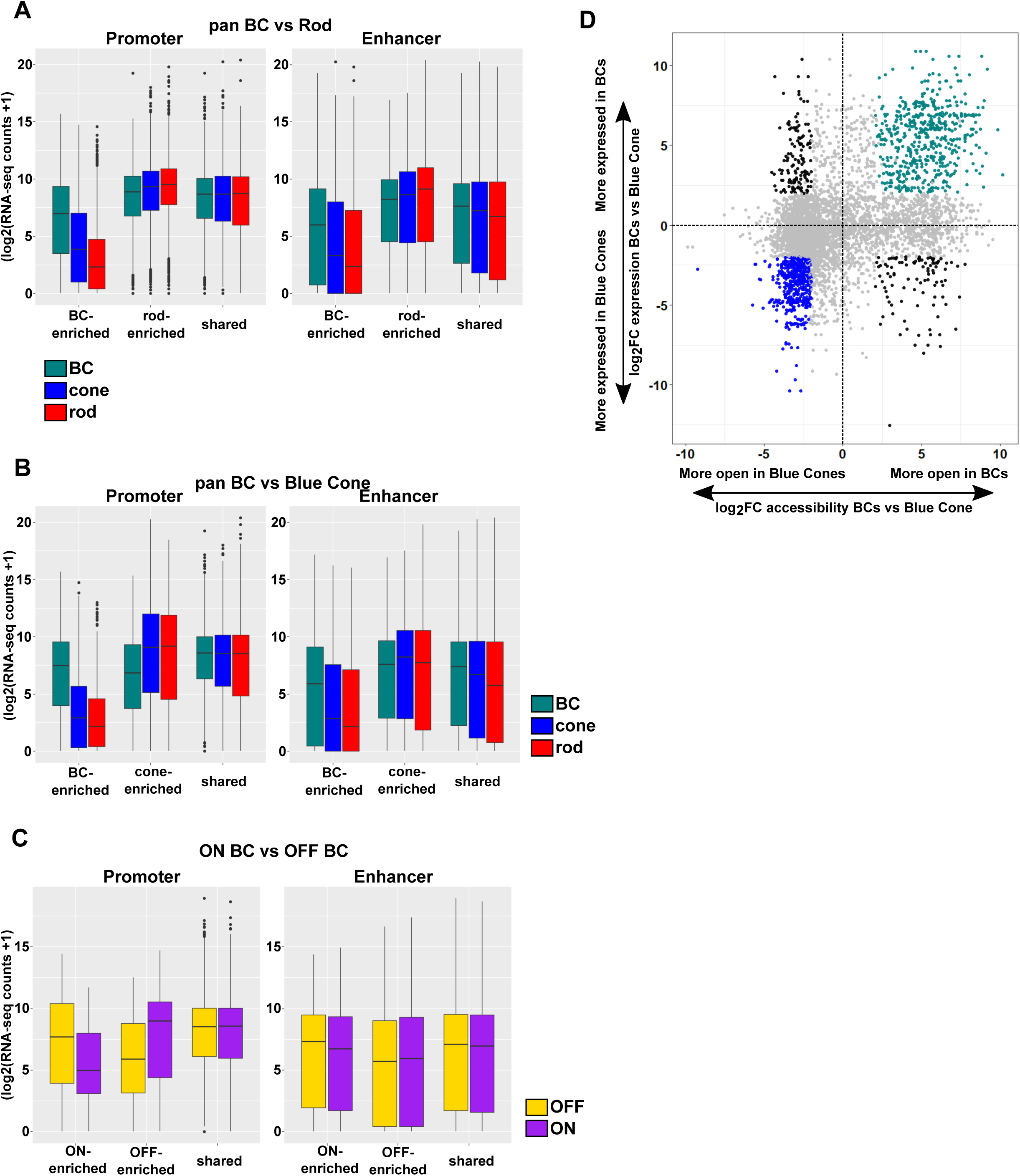
Correlation between chromatin accessibility and expression of associated genes. **A-C**. Differentially enriched ATAC-seq peaks were mapped to genes based on proximity to the nearest TSS. Peaks within −1000 bp upstream and +100 bp downstream of a TSS are defined as ‘Promoter’ peaks, whereas intergenic or intronic peaks are defined as ‘Enhancer’ peaks. Boxplots show RNA-seq expression data for genes associated with peaks that are differentially accessible in (**A**) pan BC vs rod, (**B**) pan BC vs blue cone and (**C**) ON bipolar cells vs OFF bipolar cells. (**D**). Scatterplot as in Fig. 5B showing chromatin accessibility and associated gene expression of ATAC-seq peaks identified in Fig. 5A in bipolar cells and blue cones. Peaks with four-fold greater accessibility and associated gene expression (FDR < 0.05 for both accessibility and gene expression) in bipolar cells are shown in green (n = 542), while those of blue cones are in blue (n = 505). Shared peaks and those associated with genes that have less than four-fold differences in expression are in grey. Peaks with discordant chromatin accessibility and associated gene expression (n = 234) are shown in black.

**Supplementary Figure S6.**
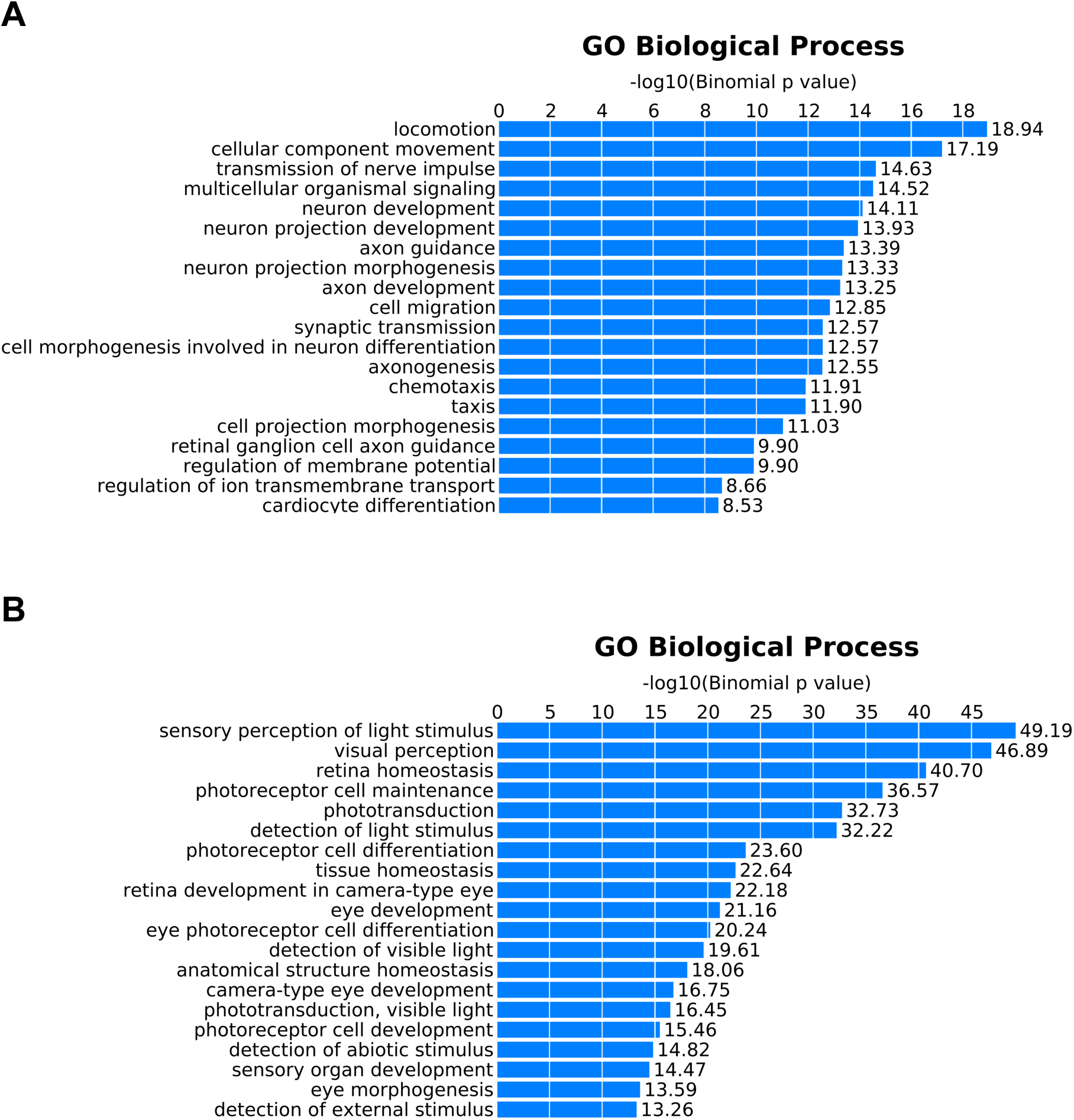
Gene Ontology (GO) annotation of peaks exhibiting correlated chromatin accessibility and associated gene expression. GO terms enriched in peaks that show positive correlations between chromatin accessibility and assigned gene expression in (A) bipolar cells or (B) photoreceptors. Peaks identified in Figure 5B and Figure S5E as bipolar-enriched (green) or photoreceptor-enriched (red or blue for rods or blue cones, respectively) were combined to create unique lists for bipolar cells (n = 833) and photoreceptors (n = 901) which were analyzed using GREAT^55^.

## Supplementary table legends

**Supplementary Table S1. Biological replicate and sequencing metrics for ATAC-seq and RNA-seq.** ‘Raw sequencing reads’ are the number of paired reads for each sample. ‘Processed reads’ are those reads remaining after filtering out those that are improperly paired, have poor mapping quality, align to the mitochondrial genome, align to ENCODE blacklisted regions, or arise from PCR duplicates. RIN = RNA integrity number.

**Supplemental Table S2. Primers used in this work.** Primers used in creation of *Gnb3* promoter constructs and in qPCR experiments are listed.

**Supplemental Table S3. Datasets and accessions.**

**Supplemental Table S4. Annotated ATAC-seq peaks and counts.** Raw count data for all ATAC-seq peaks identified in photoreceptor and bipolar cell populations. Peaks identified in individual replicates from each cell type are shown on separate sheets.

**Supplemental Table S5. Differentially expressed genes.** Genes identified as differentially expressed between aggregate bipolar cells and either rod or blue cone, and between ON and OFF bipolar cell populations are shown on separate sheets. ‘Specificity’ indicates which cell type expressed the gene more highly. For pan BC vs rod and cone, genes identified as putative transcription factors are identified by their TF family. Genes absent from the Drop-seq data shown in Figure S2 are indicated among those that are differentially expressed between ON and OFF bipolar cells.

**Supplemental Table S6. GO analysis of differentially expressed genes.** Enriched GO terms for biological processes obtained from geneontology.org. Outputs for genes enriched in photoreceptor and bipolar cells are shown on separate sheets. Input gene lists were filtered based on fold-change in expression and minimum read counts to identify those most highly enriched in photoreceptor (n=818) and bipolar cells (n=832). A list of all genes identified by RNA-seq in either cell class was used as a reference.

**Supplemental Table S7. Differentially accessible regions.** ATAC-seq peaks, normalized read counts, fold-change values, adjusted p-values and assigned genes are listed on separate sheets for each comparison. ‘Specificity’ indicates the cell type in which the peak is more highly accessible. ‘Shared-unfiltered’ peaks are those that are not differentially accessible when comparing bipolar cells versus photoreceptors (fold-change values <2 and >-2). ‘Retina’ peaks are those shown in Figure 5A, and have been filtered to remove those accessible in B cell, brain and liver. Peaks with correlated gene expression identified in Figure 5C and S5D are indicated

**Supplemental Table S8. Known motifs enriched in enhancers of bipolar cell populations.** Enrichment of all 319 motifs in the HOMER database for aggregate, ON-, and OFF-bipolar cells, each on separate sheets. A comparison of the proportional enrichment for each motif between aggregate bipolar cells and rod, blue and green cones is included on a separate sheet. A complete list of sequence logos and position weight matrices for individual motifs is available online in the HOMER motif database: http://homer.salk.edu/homer/motif/HomerMotifDB/homerResults.html.

## Acknowledgements

The authors would like to thank Leo Volkov and Yohey Ogawa for critical reading of the manuscript. The *Otx2*-GFP mouse line was a generous gift from Dr. Thomas Lamonerie (Université Côte d’Azur), and the *Grm6*-YFP line was a generous gift from Dr. Daniel Kerschensteiner (Washington University). We also credit the ENCODE consortium for the DNAse-seq datasets, the Genome Technology Access Core (GTAC) in the Department of Genetics at Washington University in St. Louis for next-generation sequencing, and the Flow Cytometry Core in the Department of Pathology and Immunology at Washington University in Saint Louis for FACS services. This work was supported by the National Institutes of Health (EY025196, EY026672, and EY024958 to J.C.C. and T32EY013360 and F32EY029571 to D.P.M.)

## Materials and Methods

### Mouse models

All animal experiments were carried out in accordance with the regulations of the IACUC at Washington University in St. Louis. Retinal dissociation and FAC sorting were carried out using *Otx2*-GFP or *Otx2*-GFP; *Grm6*-YFP mice. All *Otx2*-GFP mice were heterozygous for a GFP cassette inserted at the C-terminus of the endogenous *Otx2* locus^37^. The Grm6-YFP line harbors a YFP transgene driven by the *Grm6* promoter^38^. All electroporation experiments were carried out in CD1 mice.

### Retinal dissociation and FACS

Following dissection, retinas of 6-8 week old mice were dissociated with papain as described previously^67^. Briefly, two retinas were incubated in 400 µl of calcium/magnesium free Hanks’ Balanced Salt Solution (HBSS) (Thermo Fisher) containing 0.65 mg papain (Worthington Biochem) for 10 min at 37°C. Cells were then washed in a DMEM (Thermo Fisher) solution containing100 units DNAse1 (Roche) and incubated an additional 5 minutes at 37°C. Cells were then resuspended in 600µl of sorting buffer (2.5 mM EDTA, 25 mM HEPES, 1% BSA in HBSS) and used directly for sorting. Cells were sorted on a FACS Aria-II (BD biosciences) with gates based on forward scatter, side scatter, and GFP fluorescence. OFF bipolar cell populations (*Otx2*-GFP^+^;*Grm6*-YFP^−^) were immediately sorted a second time to increase purity.

### Generation of reporter constructs

An 820 bp region encompassing part of the *Gnb3* 5’ UTR and upstream sequence was amplified from mouse genomic DNA. Site-directed mutagenesis by overlap extension was used to modify K50 sites^68^. The resultant PCR products were digested and ligated into GFP or DsRed reporter vectors derived from pCAGGS^69^. After verification by sequencing, plasmid DNA was resuspended in PBS at a concentration of ~6-7µg/µl prior to injection. All primers are listed in Table S2.

### *In-vivo* and explant electroporation

*In-vivo* subretinal injection and electroporation of newborn CD1 mice was performed as previously described^70^. Briefly, mice were first anesthetized on ice. A 30-gauge needle was then used to incise the eyelid and puncture the sclera, and a Hamilton syringe with a 33-gauge blunt-tipped needle was used to inject the DNA into the subretinal space. Tweezer electrodes placed across the head were then used to electroporate with 5 square pulses of 80 volts and 50 millisecond duration at 950 millisecond intervals. Explant electroporation was carried out as described previously^69^, except that the electroporation chamber contained a solution of 0.5µg/µl DNA in PBS.

### Retinal tissue sectioning and imaging

Eyes were removed at P21, punctured with a 26-gauge needle, and incubated in 4% paraformaldehyde for 5 minutes before dissection to remove the cornea. The lens was removed, and eyes were then incubated for an additional 45 minutes in 4% paraformaldehyde. Eye cups were next washed in PBS and incubated overnight at 4°C in 30% sucrose-PBS. The following day, eye cups were incubated in a 1:1 mixture of OCT compound (Sakura) and sucrose-PBS before being flash frozen in OCT and stored at −80°C. Retinal sections of 14 µm were cut using a cryostat (Leica CryoCut 1800), mounted on Superfrost Plus slides (Fisher), and stored at −20°C. Prior to placement of cover slips, slides were washed with PBS to remove OCT. The sections were then stained with DAPI, and coverslips were mounted using Vectashield mounting medium (Vectorlabs). Retinal sections were imaged on a Zeiss 880 laser-scanning confocal microscopy in the Washington University Center for Cellular Imaging (WUCCI) Core.

### ATAC-seq

Transposition and library preparation from sorted cell populations were performed as previously described^71^. Briefly, 30,000-100,000 sorted cells were pelleted at 500 G and washed twice in ice-cold PBS before lysis. Transposition reactions were incubated at 37°C for 30 minutes and purified using a Qiagen MiniElute PCR Purification kit. Libraries were amplified with Phusion High-Fidelity DNA Polymerase (NEB). Cycle number was calibrated by a parallel qPCR reaction. Gel electrophoresis was used to assess library quality, and final libraries were quantified using KAPA Library Quantification Kit (KAPA Biosystems). Equimolar concentrations of each library were pooled and run on an Illumina HiSeq2500 to obtain 50 bp paired-end reads.

### ATAC-seq, DNAse-seq, and RNA-seq data processing

ATAC-seq and RNA-seq reads from bipolar cell populations were processed in an identical manner to those previously obtained from rod and cone photoreceptor cells^30^. ATAC-seq reads were aligned to the GRCm38/mm10 mouse genome assembly using Bowtie2 (v2.3.5) with a max fragment size of 2000^72^. Alignments were filtered using SAMtools (v1.9)^73^, PCR duplicates were removed using Picard (v2.19.0) (https://broadinstitute.github.io/picard/), and nucleosome-free reads were selected by removing alignments with an insertion size greater than 150 bp. Peaks were called using MACS2 (v2.1.1)^74^ and annotated with HOMER (v4.8)^47^. DNAse-seq datasets generated by ENCODE^46^ were downloaded as FASTQ files from https://www.encodeproject.org/ and processed in the same manner as ATAC-seq data. RNA-seq reads were aligned to the GRCm38/mm10 using STAR (v2.7.0d)^75^, with an index prepared for 50 base-pair reads and the RefSeq gene model, and read counts were calculated using HTSeq (v1.9)^76^. All datasets are listed in Table S3.

### Transcription factor binding site motif analysis

Motif enrichment, co-enrichment, and spacing analyses for ATAC-seq and DNAse-seq datasets were performed as described previously using HOMER (v4.8)^30,47^. Differential motif enrichment was determined using a test of equal proportions (R stats v3.5.3) to compare each motif between pan-BC and rods, blue cones or green cones. The top motifs across the three comparisons were manually filtered for redundancy and are shown in Figure 4. Motif co-occurrence analysis was performed using a list of 66 non-redundant motifs^30^ to which the motif for LHX3 (representing a Q50 HD motif) was manually added. For purposes of the analysis, peaks from rod, green cones and blue cones were merged to obtain a ‘photoreceptor’ peak list, while those from pan-, ON- and OFF-bipolar cells were merged to create a ‘bipolar cell’ peak list. Enrichment for co-occurrence was calculated by taking the log_2_(observed pairs/expected pairs). Expected frequency of individual pairs was estimated from the frequency of each motif within the pair (peaks with motif 1 × peaks with motif 2 ÷ number of total peaks). Motif spacing was analyzed for the top enriched peaks shown in Figure 4. The same set of peaks for co-occurrence were centered on individual K50 or Q50 motifs, and the density of flanking secondary motifs was plotted on either strand.

### Identification of differentially accessible peaks and differentially expressed genes

DESeq2 (v1.14.1)^77^ was used to test for differential expression or differential accessibility using a log_2_ fold-change threshold of 1 and an FDR of 0.05. For comparison of ATAC-seq data with DNAse-seq data from non-retinal tissues (Fig. 5), photoreceptors were collapsed into a single level. Differentially expressed genes are listed in Table S5, and differentially accessible regions are listed in Table S7. For each comparison (i.e. ON versus OFF, pan-BC versus rod), gene expression stemming from low-level contamination of bipolar cell populations with either rod or cone photoreceptors was filtered out. When comparing pan-BC to photoreceptor populations, potential contaminating genes from the alternate photoreceptor type (i.e. rod genes identified as enriched in pan-BC versus blue cone) were identified as those highly expressed (>16-fold) and specific to rod compared to blue cone, and also more highly expressed in rod compared to pan-BC (at least four-fold). In comparing ON and OFF bipolar cells, genes enriched in each bipolar cell population were filtered for those which were also identified as highly specific to either photoreceptor population compared to the enriched bipolar cell type. For example, genes increased in OFF-compared to ON-bipolar cells were filtered for genes that were also highly enriched (>16-fold) in rods and blue cones compared to OFF bipolar cells. In total, 38 genes were filtered from those identified as enriched in pan-BC compared to photoreceptors, and 39 genes were filtered from the ON versus OFF comparison (12 from the ON-enriched, 27 from the OFF-enriched). These genes include those known to be expressed at very high levels in either rod or cone photoreceptors^78^.

### RNA isolation, qPCR and RNA-seq

Sorted bipolar cell populations were resuspended in 500 µl TRIzol reagent (Invitrogen), and RNA was isolated according to manufacturer’s instructions. Prior to sequencing, RNA quality was analyzed using an Agilent Bioanalyzer. cDNA was prepared using the SMARTer Ultra Low RNA kit for Illumina Sequencing-HV (Clontech) per manufacturer’s instructions. cDNA was fragmented using a Covaris E210 sonicator using duty cycle 10, intensity 5, cycles/burst 200, time 180 seconds. cDNA was blunt-ended, an ‘A’ base added to the 3′ ends, and Illumina sequencing adapters were ligated to the ends of the cDNAs. Ligated fragments were amplified for 12 cycles using primers incorporating unique index tags. Replicate libraries from each bipolar cell population were pooled in equimolar ratios and sequenced on an Illumina HiSeq 3000 (single-end 50 bp reads). For qPCR, RNA samples were treated with TURBO DNAse (Invitrogen) and cDNA was synthesized with SuperScript IV (Invitrogen) and oligo(dT) primers according to manufacturer’s instructions. Expression was normalized to the average of reference genes *Gapdh*, *Sdha*, *Hprt*, and *Pgk1*. Primers for *Grm6*, *Gnat1*, *Lhx1*, *Pax6*, *Rlbp1*, *Slc17a6*, *Vsx2*, *Grik1*, and *Tacr3* are listed in Table S2.

